# Effect of RNA m^6^A methyltransferase activation by a low molecular weight compound on anxiety- and depression-related behaviours, monoamine neurochemistry and striatal gene expression in the rat

**DOI:** 10.1101/2024.01.08.574593

**Authors:** Margus Kanarik, Kristi Liiver, Marianna Školnaja, Indrek Teino, Tõnis Org, Karita Laugus, Ruth Shimmo, Mati Karelson, Mart Saarma, Jaanus Harro

## Abstract

Modification of mRNA by methylation is involved in post-transcriptional regulation of gene expression by affecting the splicing, transport, stability and translation of mRNA. Methylation of adenosine at N^6^ (m^6^A) is the most common and most important cellular modification occurring in the mRNA of eukaryotes. Evidence that m^6^A mRNA methylation is involved in regulation of stress response and that its dysregulation may contribute to the pathogenesis of neuropsychiatric disorders is accumulating. We have examined the acute and subchronic (up to 18 days once per day intraperitoneally) effect of the first METTL3/METTL14 activator compound CHMA1004 (methyl-piperazine-2-carboxylate) at two doses (1 and 5 mg/kg) in male and female rats. CHMA1004 had a profound locomotor activating and anxiolytic-like profile in open field and elevated zero-maze tests. In female rats sucrose consumption and swimming in Porsolt’s test were increased. Nevertheless, CHMA1004 did not exhibit strong psychostimulant-like properties: CHMA1004 had no effect on 50-kHz ultrasonic vocalizations except that it reduced the baseline difference between male and female animals, and acute drug treatment had no effect on extracellular dopamine levels in striatum. Subchronic CHMA1004 altered *ex vivo* catecholamine levels in several brain regions. RNA sequencing of female rat striata after subchronic CHMA1004 treatment revealed changes in the expression of a number of genes linked to dopamine neuron viability, neurodegeneration, depression, anxiety and stress response. Conclusively, the first-in-class METTL3/METTL14 activator compound CHMA1004 increased locomotor activity and elicited anxiolytic-like effects after systemic administration, demonstrating tha pharmacological activation of RNA m^6^A methylation has potential for neuropsychiatric drug development.

## Introduction

Psychiatric disorders mostly develop if genetic vulnerability meets life adversities. These disorders often have chronic and relapsing course. Triggering of the pathogenetic cascade by life events and maintenance of the new meta-stable state of the CNS is likely to be mediated by multiple epigenetic and downstream regulatory mechanisms. Epigenetic DNA modifications, especially DNA methylation, in psychiatric conditions such as affective, anxiety and substance use disorders or aggressiveness have garnered much scrutiny in recent years (Juruena et al., 2021; Panariello et al., 2022; Harvanek et al., 2023; Persaud and Cates, 2023; Pishva et al., 2023). However, limited attention has been paid to modification of mRNA by methylation. mRNA methylation is involved in post-transcriptional regulation of gene expression by affecting the stability and translation of mRNA. Methylation of adenosine at N^6^ (m^6^A) is the most common and most important cellular modification occurring in the mRNA of eukaryotes. m^6^A methylation regulates splicing, transport, stability and translation of mRNA (Zaccara et al., 2019). Three main classes of proteins regulate these processes. m^6^A formation is catalysed by a methyltransferase complex containing methyltransferase-like 3 (METTL3), methyltransferase-like 14 (METTL14), and Wilms-tumor-1-associated protein (WTAP), and by a single subunit enzyme methyltransferase-like 16 (METTL16). RNA m^6^A demethylation is catalysed by the fat mass and obesity-associated protein FTO and the RNA demethylase enzyme ALKBH5. These enzymes modify usually only one but sometimes up to three adenosine residues in the mRNA molecules generally located in the 3′untranslated region. RNA sequences containing m^6^A residues are recognized by a dozen of RNA binding proteins, e.g., YTHDC1, YTHDC2, YTHDF1, YTHDF2 and YTHDF3 that regulate the fate of mRNA (Meyer and Jaffrey, 2017). m^6^A modification occurs in all organs but is particularly abundant in mammalian brain. There is increasing evidence on how mRNA m^6^A methylation is involved in regulation of behaviour, suggestive that its dysregulation may contribute to the development of and pathogenesis of neuropsychiatric disorders (Zaccara et al. 2019; Livneh et al., 2020; Selberg et al., 2021; Shafik et al., 2022; Su et al., 2023).

A pioneering study demonstrated that interference with RNA m^6^A methylation affects clock gene transcription and by this means the length of circadian period in mice (Fustin et al., 2013). More recently, artificial lighting was shown to affect RNA m^6^A methylation and corresponding gene expression in the rat lateral habenula (Li Y et al., 2023), a brain region pivotal for fronto-limbic to brainstem connections and implicated in affective and substance use disorders (Ables et al., 2023). Recent studies have suggested the involvement of cerebral RNA m^6^A methylation in stress response, whereas m^6^A and its regulatory gene expression levels can change in a regionally specific manner (Engel et al., 2018). In the mouse prefrontal cortex m^6^A methylation was increased in response to submission to behavioural tests, especially prominently after fear conditioning, and if Fto was knocked down, cued fear memory was increased (Widagdo et al., 2016). Contextual fear memory was also increased after viral expression of shRNA targeted against Fto in dorsal hippocampus (Walters et al., 2017). Consolidation of long-term memory was found to be dependent on METTL3 function (Zhang et al., 2018). Another study reported a positive correlation of m^6^A methylation and the methyltransferase *Mettl3* levels with ability of contextual fear discrimination and demonstrated that expression of *Mettl3* in hippocampal glutamatergic neurons is crucial for this type of learning and memory (Chang et al., 2022). In learned helpless rats the expression of several so-called plasticity genes in hippocampus was downregulated along with enrichment in their m^6^A methylation (Roy et al., 2022). In summary, RNA m^6^A methylation serves as a crucial regulation mechanism at the epitranscriptomic level to shape synaptic plasticity and behaviour (Malovic and Pandey, 2023) and involvement of RNA m^6^A methylation appears in multiple behavioural conditions relevant to anxiety, but so far evidence for a role of m^6^A methylation has remained correlational.

*In silico*-based approach that departed from the crystal structure of the METTL3/METTL14 complex with S-adenosyl-L-homocysteine (Wang P et al., 2016) enabled some of us to identify the first small molecule ligands that bind to METTL3/METTL14/WTAP and function as RNA methylation activators (Selberg et al., 2019). In particular, as measured using the surface plasmon resonance, the compounds increased 200-fold the binding efficiency of the RNA methylating substrate S-adenosylmethionine (SAM) to the METTL3/METTL14/WTAP protein complex. The treatment of cells with these compounds activating the METTL3/METTL14/WTAP complex led to the increase of m^6^A modified nucleotides in the substrate RNA. This is in a sharp contrast to the suppression of the methylation by a non-specific methylation inhibitor sinefungin. The cytotoxicity screening on the number of endpoints including cell count, nuclear size, DNA structure, cell membrane permeability, mitochondrial mass, mitochondrial membrane potential and cytochrome c release did not show any aberrations up to the 200 µM concentration of compounds (acute oral toxicity LD50 > 2,000 mg/kg). The compounds do not inhibit hERG channel and the monoamine oxidase. The compounds have good water solubility (> 100 µM) and the half-lives of the best compound CHMA1004 are t1/2 = 354 min in human plasma and t1/2 = 60.5 min in rat plasma, respectively (data available on request). The computational modelling on other methyltransferases (DNA and histone methyltransferases) did not show any strong binding of the m^6^A methylation activators to these enzymes, thus they are predicted to be very specific to METTL3/METTL14/WTAP protein complex. Two of these compounds were recently found to support the survival of mouse dopamine neurons in growth factor deprivation assay. and to protect them from 6-OHDA toxicity. Importantly, CHMA1004 (methyl-piperazine-2- carboxylate) protected stem cell-derived human dopamine neurons against 6-OHDA induced cell death. Furthermore, intrastriatal administration of CHMA1004 attenuated amphetamine-induced ipsilateral rotations after 6-OHDA treatment and increased dopamine neuron fibre density in the rat model of PD (Yu et al., 2023). Given the massive arborization of dopaminergic axons (Matsuda et al., 2009), dopaminergic neural projections may be particularly sensitive to the epitranscriptomic regulation of mRNA translation that occurs in the axonal and dendritic ribosomes (Biever et al., 2020).

Thus, taking into account the indirect evidence reviewed above on the role of RNA m^6^A methylation in anxiety and depression, and the implication of dopaminergic neurotransmission in these disorders that often are co-morbid with addictions (Rehm et al., 2016), we aimed to examine the effect of the novel METTL3/METTL14 activator compound CHMA1004 in a battery of anxiolytic and antidepressant screening tests in the rat, its effect on emission of ultrasonic vocalizations and effects of acute amphetamine treatment. The effect of subchronic administration of CHMA1004 on monoamine neurochemistry in several brain regions and on gene expression in striatum, and its acute effect on dopamine overflow, were also examined. We found that systemic delivery of METTL3/METTL14 activator compound CHMA1004 to rats elicited an increase in locomotor activity and attenuation of anxiety- and depression-related behaviours.

## Materials and Methods

### Animals

In the behavioural experiments, twenty seven Wistar rats (15 female and 12 male; 3 months of age at the beginning of the experiment; body weight 290±4 g and 499±13 g respectively; bred locally, parent rats obtained from Charles River) were given CHMA1004 (1 mg/kg or 5 mg/kg) dissolved in saline intraperitoneally (IP; 1 ml/kg) once in a day. Control group received vehicle (saline). Each group consisted of 5 female and 4 male rats. In the microdialysis experiment 19 male and 16 female rats with the same background were used; their allocation to treatment groups is given in the Microdialysis section. Animals were housed 4-5 per cage in standard transparent polypropylene cages under controlled light cycle (lights on 08:00-20:00) and temperature (22°-24°C), with free access to water and food pellets (v1534-000 universal maintenance diet, ssniff Spezialdiäten GmbH, Soest, Germany). The studies were approved by the Ethical Committee for Animal Experiments of the Estonian Ministry of Rural Affairs (permissions nr 127, 134, 149).

### General procedure

The procedure is illustrated on Figure 1. Thirty minutes after the first IP administration of CHMA1004 or vehicle an open field test was conducted with recording of ultrasonic vocalizations (USVs). A series of behavioural tests were performed in the course of daily drug treatment, while all behavioural tests were started 30 min after drug administration. Open field tests were performed again after 7 and 13 days of treatment immediately after conducting elevated zero-maze tests. On Day 8 social interaction test was performed, and on Days 10 and 11, the forced swimming test. After two weeks of daily CHMA1004 treatment, locomotor activity was measured after administration of amphetamine (1 mg/kg, IP), and USVs were recorded. USVs were also recorded on Day 9 in a novel cage similar to the housing environment. On Day 15, the sucrose preference test was conducted. After 17 or 18 days of daily drug treatment rats were sacrificed 30 min after the last injection by decapitation, the brains dissected as previously described (Liiver et al., 2023) and samples immediately frozen in liquid nitrogen; selected regions were used for the measurement of monoamine levels, or RNA sequencing (striatum).

**Figure 1.**
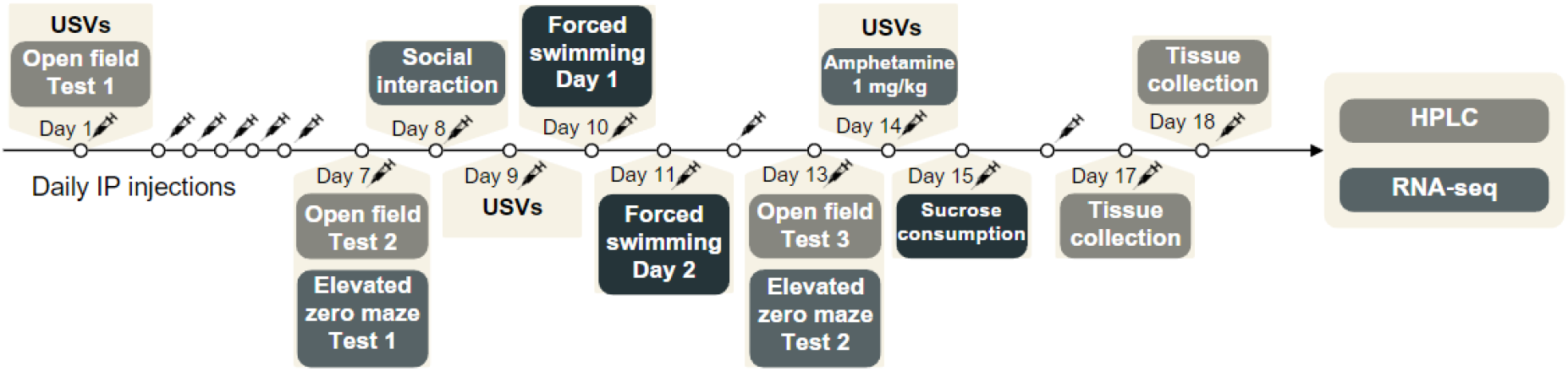
Experimental design of behavioural and *ex vivo* biochemical studies of the effect of acute and subchronic treatment with CHMA1004. HPLC, high performance liquid chromatography; IP, intraperitoneal; RNA-seq, RNA sequencing; USVs, recording of ultrasonic vocalization.

All procedures, except the sucrose preference test, occurred during the lights-on phase. Animals were transported 30 min before the start of testing to the room of behavioural experiments to habituate, and tested (within the sex group) in randomized order. All animal experiments were in compliance with the ARRIVE guidelines and were carried out in accordance with EU Directive 2010/63/EU for animal experiments; the experimental protocol was approved by the Animal Experimentation Committee at the Estonian Ministry of Rural Affairs.

### Open field test

Open field test was carried out in a 0.5 x 1.0 m metal box with side walls 40 cm high (Tõnissaar et al., 2008). The area was divided into eight squares of equal size. Time spent moving around was recorded, and square crossings and rearing in 4 min were counted, as was the number of faecal boli left on the open field. The apparatus was cleaned with laboratory tissue dampened with water after each animal.

### Ultrasonic vocalizations

The microphone of an ultrasound recording device (Avisoft Ultra Sound Gate 116–200, Avisoft Bioacoustics, Berlin, Germany) was located about 30 cm from the floor of the apparatus, recording USVs with a sampling rate of 300 kHz in 16-bit format on a computer hard drive (Kõiv et al., 2021). The files were recorded with Avisoft-RECORDER software and later analysed with Avisoft SASLab Pro (Avisoft Bioacoustics, Berlin, Germany) software. Spectrograms were created using the Fast Fourier Transform algorithm (1024 FFT length, 75% frame, Hamming window, and 75% time window overlap). USVs were manually marked and saved on the spectrogram by an observer blind to experimental conditions. Fifty kHz USVs with frequencies over 35 kHz, including flat, frequency-modulated and trill calls) and 22 kHz (<35 kHz) USVs were counted.

### Elevated zero-maze

The elevated zero-maze test (Shepherd et al., 1994) was conducted as previously described (Matto et al., 1997). An elevated annular platform (outer Ø 105 cm, width 10 cm, 72 cm above the floor) was equally divided into two opposing enclosed quadrants (height of the walls 28 cm) that were connected by open quadrants (height of the edge 1 cm, 5-11 lux in the open part). The open quadrants were also divided into three equidistant parts to quantify the locomotor activity. The rat was placed at the outset of one of the closed quadrants and was observed at a distance for 5 min. The measures taken were stretched-attend postures toward the open quadrants, latency to enter the open quadrant with all four paws (s), entries into the open quadrants, and time spent (s) in the open quadrants. The apparatus was cleaned with moist tissue after every rat.

### Social interaction test

The social interaction test introduced by Sandra File (reviewed by File and Hyde, 1978) was conducted as described previously (Tõnissaar et al., 2004) with the variance that animals were not single-housed. Two unfamiliar, weight-matched rats were placed in the opposite corners of a brightly lit chamber (30×30×60 cm) with floor covered with wood shavings. The behaviour of the animals was concurrently observed for 10 min and the total time spent in active social behaviour (allogrooming, sniffing the partner, crawling under and over, following) was recorded.

### Forced swimming test

The forced swimming test (Porsolt et al., 1978) was carried out as described previously (Mällo et al., 2009). On the first day of the experiment, the procedure lasted 15 min and the re-exposure 24 h later lasted 5 min. The measurements were based on the behavioural categories described by Armario et al. (1988); the first 5-min period of the first day was counted. A rat was judged to be immobile when it remained floating in the water with all limbs motionless or made minimal movements in order to maintain its head above the water, and to struggle whenever it made intense movements of all the four limbs with the two front paws breaking the surface of the water or touching the walls of the tank. The time spent in swimming represents the rat making active swimming motions, more than necessary to merely maintain its head above water.

### Amphetamine treatment effect

D-Amphetamine (1 mg/kg IP) was dissolved in saline and administered 15 min prior the animal was placed in a standard group-housing cage (55×33 cm) for assessment of stimulant-induced locomotor response and ultrasonic vocalisation (Kõiv et al., 2016). Horizontal locomotion was assessed from a 12-minute video by counting line crossings with four paws of the boundaries of six equal squares on the cage floor, rearings were counted when the animal stood on hind legs with the body at least at a 45° angle. Data were analysed by 2-min bins.

### Locomotion and ultrasound vocalizations in a novel home-cage like environment

Measurement of USVs and spontaneous horizontal and vertical locomotion was conducted in a standard novel home cage like box (for eight minutes) with a similar protocol as previously described for amphetamine induced behavioural measurements.

### Measurement of sucrose intake and preference

The sucrose preference test was carried out during the dark phase with no previous food or water deprivation (Kõiv et al., 2019). The animals were placed into single cages 1 hour before the sucrose preference testing and group-housed again after that. Food was available during sucrose preference test. Two bottles were used, one filled with 1% sucrose solution and the other with water. Liquid consumption was measured by weighing previously weighed sucrose-solution and water bottles 1 and 12 h after the beginning of the sucrose preference test. Sucrose preference was calculated as the proportion of sucrose solution consumption out of the total consumption of water and sucrose solution.

### High performance liquid chromatography (HPLC) in ex vivo studies

Monoamines and their metabolites were assayed by HPLC with electrochemical (amperometric) detection as described previously (Kõiv et al., 2019). Rat brain tissues were homogenised with an ultrasonic homogeniser (Bandelin Sonopuls, Bandelin Electronic, Berlin, Germany) in ice-cold solution of 0.1 M perchloric acid containing 5 mM of sodium bisulfite and 0.4 mM EDTA to avoid oxidation. The homogenate was then centrifuged at 14,000 rpm for 10 min at 4°C. Aliquots (10 μl) of the supernatant obtained were chromatographed on a Luna C18(2) column (150×2 mm, 5 μm). The separation was done in isocratic elution mode at column temperature of 30°C, using the mobile phase containing 0.05 M sodium citrate buffer at pH 3.7, 0.02 mM EDTA, 1 mM KCl, 1 mM sodium octanesulphonate and 7.5% acetonitrile. The two used chromatography systems consisted of an isocratic pump (HP1100, Agilent, Waldbronn, Germany or Shimadzu LC-20AD, Japan), a temperature-regulated autosampler (HP1100 or Shimadzu SIL-20AC), a temperature-regulated column compartment and an HP 1049 electrochemical detector (Agilent, Waldbronn, Germany) with a glassy carbon electrode (HP flow cell or Antec VT-03 flow cell). The measurements were done at an electrode potential of +0.7 V versus the Ag/AgCl reference electrode. The limits of detection at signal-to-noise ratio=3 were as follows (expressed as pmol/mg tissue for each): 0.08 for dopamine (DA), 0.10 for homovanillic acid (HVA), 0.05 for 3,4-dihydroxyphenylacetic acid (DOPAC), 0.08 for serotonin (5-HT), 0.04 for 5-hydroxyindoleacetic acid (5-HIAA), 0.07 for noradrenaline (NA), 0.03 for normetanephrine (NMN), and 0.01 for 3-methoxytyramine (3-MT).

### RNA extraction and mRNA sequencing

RNA was extracted from female rat striatal samples using FavorPrep^TM^ Tissue Total RNA Mini Kit (FATRK 001-2, Favorgen, Taiwan) according to the manufacturer’s protocol. Briefly, the FARB lysis buffer, containing ß-mercaptoethanol, was added to the tissue sample, grinding was performed with a micropestle followed by homogenization through a 20-G needle syringe. The lysate was cleared by using the filter column followed by addition of one volume of 70% ethanol. The sample was then transferred to the FARB Mini Column, washed and dried by centrifugation at 18,000 g for three minutes. Total RNA was eluted in 30 µl of nuclease-free water, concentration was measured by NanoDrop 1000 spectrophotometer (ThermoFisher Scientific Inc., Waltham, MA, USA) and stored at −80°C.

Sequencing was performed by Novogene Co., Ltd. (Beijing, China). Quality control and preprocessing of the fastq files, containing raw reads from Illumina 150bp paired-end sequencing, was conducted with fastp v0.23.2 using default parameters (Chen et al., 2018). Strand specific alignment to the rat rn7 genome was carried out using HISAT2 v2.2.1. (paired end, Strand -FR). FeatureCounts v1.6.4 (Liao et al., 2014) was used for transcript counting using *Rattus_norvegicus*.mRatBN7.2.108.gtf annotation from Ensembl database as a reference (Martin et al., 2023). Differential gene expression between pairwise groups was performed with DeSeq2 v22.11.40.6 (Love et al., 2014) using poscounts method for size factor estimation. Genes with p-value smaller than 0.05 were considered differentially expressed. SynGO portal was used for ID conversion (Koopmans et al., 2019). GO enrichment analysis was performed with DAVID 2021 (Sherman et al., 2022). Two samples were excluded from the subsequent analysis because of low alignment rate (<5%) to the reference rn7 genome, and one sample was considered as an outlier by visual inspection of PCA plot and omitted from subsequent analysis; consequently, further analysis included 4 rats per group. Volcano plot was created using R ggplot2 based tool in Galaxy (Version 0.0.3) using output from DESeq2.

### Microdialysis experiments

*In vivo* microdialysis was carried out to assess the effect of acute treatment with CHMA1004 on extracellular levels of dopamine in striatum, essentially as previously described [O’Leary et al., 2016]. The animals were anaesthetized with ketamine/medetomide anaesthesia (60 and 0.5 mg/kg IP, respectively) and mounted in a Stoelting stereotactic frame. A self-made concentric Y-shaped microdialysis probe with 7 mm shaft length and 3 mm active tip was implanted into the left dorsal striatum according to the following coordinates: AP +0.7; ML +3.0; DV –7.0, according to Paxinos and Watson (1986). The dialysis membrane used was polyacrylonitrile/sodium methalyl sulphonate copolymer (Filtral 12; i.d.: 0.22 mm; o.d.: 0.31 mm; AN 69, Hospal, Bologna, Italy). Two stainless steel screws and dental cement were used to fix the probe to the scull. After the surgery, anaesthesia was reversed by administration of atipamezole (1 mg/kg, SC) and meloxicam (1-2 mg/kg, SC) was administered for post- operative analgesia. The rats were placed in 21 cm × 36 cm × 18 cm individual cages in which they remained throughout the microdialysis experiment. Rats were given about 24 h for recovery and microdialysis procedure was conducted in awake freely moving animals. The microdialysis probe was connected via FEP tubing (i.d. 0.12 mm, AgnTho’s AB, Lidingö, Sweden) to a syringe pump (SP101, World Precision Instruments, Inc., Sarasota, FL, USA) and to a refrigerated microsampler (Univentor 820, Univentor Limited, BLB029 Bulebel Industrial Estate, Zejtun ZTN 3000, MALTA) and perfused with Ringer solution (147 mM NaCl, 4 mM KCl, 1.2 mM CaCl2, 1.0 mM MgCl2, 1.0 mM Na2HPO4; pH 7.20–7.22) at a constant rate of 1.5 μl/min. After connecting the animal to the microdialysis system, the perfusate was discarded during the first 60 min to allow stabilization. Six baseline samples were collected, followed by an i.p. injection of either saline (male rats; n=2), CHMA1004 1 mg/kg (male, n=6; female, n=6), CHMA1004 5 mg/kg (male, n=6; female, n=6) or amphetamine 0.5 mg/kg (male, n=4; female, n=4), after which another 12 samples were collected. All samples were collected with 15-min intervals. Sample vials were prefilled with 7.5 μl of 0.02 M acetic acid to prevent oxidation of dopamine. The quantity of dopamine and serotonin in the microdialysis samples was determined by high performance liquid chromatography with electrochemical detection. The chromatography system consisted of a Shimadzu LC-10AD pump and CBM-20A controller (Shimadzu Corporation, Kyoto, Japan), a Luna C18(2) 5 μm column (150 x 2 mm) kept at 30°C and Decade II digital electrochemical amperometric detector (Antec Leyden BV, the Netherlands) with electrochemical flow cell VT-03 (2 mm GC WE, ISAAC reference electrode, Antec Leyden BV, the Netherlands). The mobile phase consisted of 0.05 M sodium citrate buffered to pH 5.3, 2 mM KCl, 0.02 mM EDTA, 3.5 mM sodium octyl sulfonate and 14% acetonitrile. The mobile phase was pumped through the column at a rate of 0.2 ml/min. 5-HT and DA eluted from the column were measured with a glassy carbon working electrode maintained at a potential of +0.4 V versus Ag/AgCl reference electrode. Data were acquired using a Shimadzu LC Solution system.

### Data analysis

Behavioural and neurochemical data were first analysed by two-factor ANOVA (Sex and Treatment) with repeated measures (Test) if appropriate. Because the apparent dose-dependency of the effect of CHMA1004 was in some tests different between sexes, this was followed by one-way ANOVA (CHMA1004) with male and female dosage group pooled or separate if the interaction term was closely missing p<0.05. Subsequent pairwise comparisons were made by Fisher’s least significant difference (LSD) test. Significance threshold was set at 0.05. Statistical analysis was made using the IBM SPSS Statistics 26 software.

## Results

### Body weight and weight of adrenals

Body weight gain was similar in all treatment groups. After the 18-day treatment with CHMA1004, no statistically significant effect was found on adrenal weight (data not shown).

### Effect of acute and repeated administration of CHMA1004 on open field behaviour

Acute treatment with CHMA1004 increased time spent in active movement on the open field (Figure 2); in female rats this effect was statistically significant at 1 mg/kg and still observable after repeated treatment (Figure 3) despite of being smaller as the activity became lower (Treatment × Sex F2,21=3.49; p<0.05; Test F2,42=33.5; p<0.001). This increase of activity was statistically significant only in female rats while at the dose of 5 mg/kg a similar tendency appeared in males. Similar group differences were reflected in rearings (Treatment F2,21=4.27; p<0.05; Sex F1,21=7.96; p<0.05; Treatment × Sex F1,21=3,42; p=0.052; Test F2,42=10.3; p<0.001), and in the number of line crossings in which case the tendency of increase in males at the dose of 5 mg/kg was the closest to statistical significance (Treatment F2,21=3.37; p=0.054; Sex F1,21=6.88; p<0.05; Treatment × Sex F2,21=4.59; p<0.05; Test F2,42=11.3; p<0.001). The number of faecal boli left on the open field at the first test, a rather sensitive anxiety measure, was larger in male rats (Figure 2), and this was reduced by CHMA1004 treatment (Sex F1,21=21.4; p<0.001; Treatment F1,21=3.46; p=0.05; Treatment × Sex F2,42=4.78; p<0.05). Thus, CHMA1004 treatment elicited anxiolytic- or psychostimulant-like locomotor activation, especially in female rats, and reduced emotionality in male rats as expressed in mitigation of defecation.

**Figure 2.**
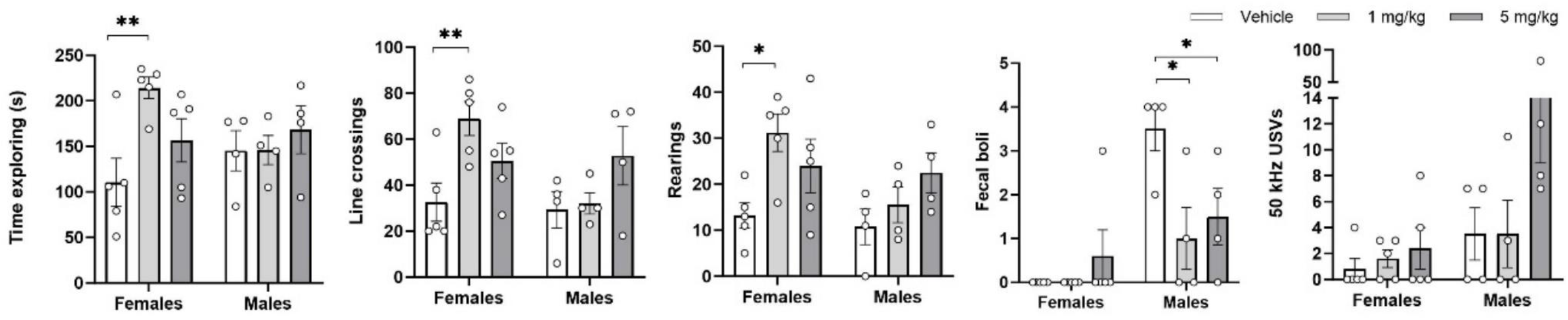
Effect of acute administration of CHMA1004 on locomotion, defecation and emission of ultrasonic vocalizations in an open field test. *p<0.05; **p<0.01 vs. vehicle. Post hoc test on rearings in male rats CHMA1004 (5 mg/kg) p=0.069.

**Figure 3.**
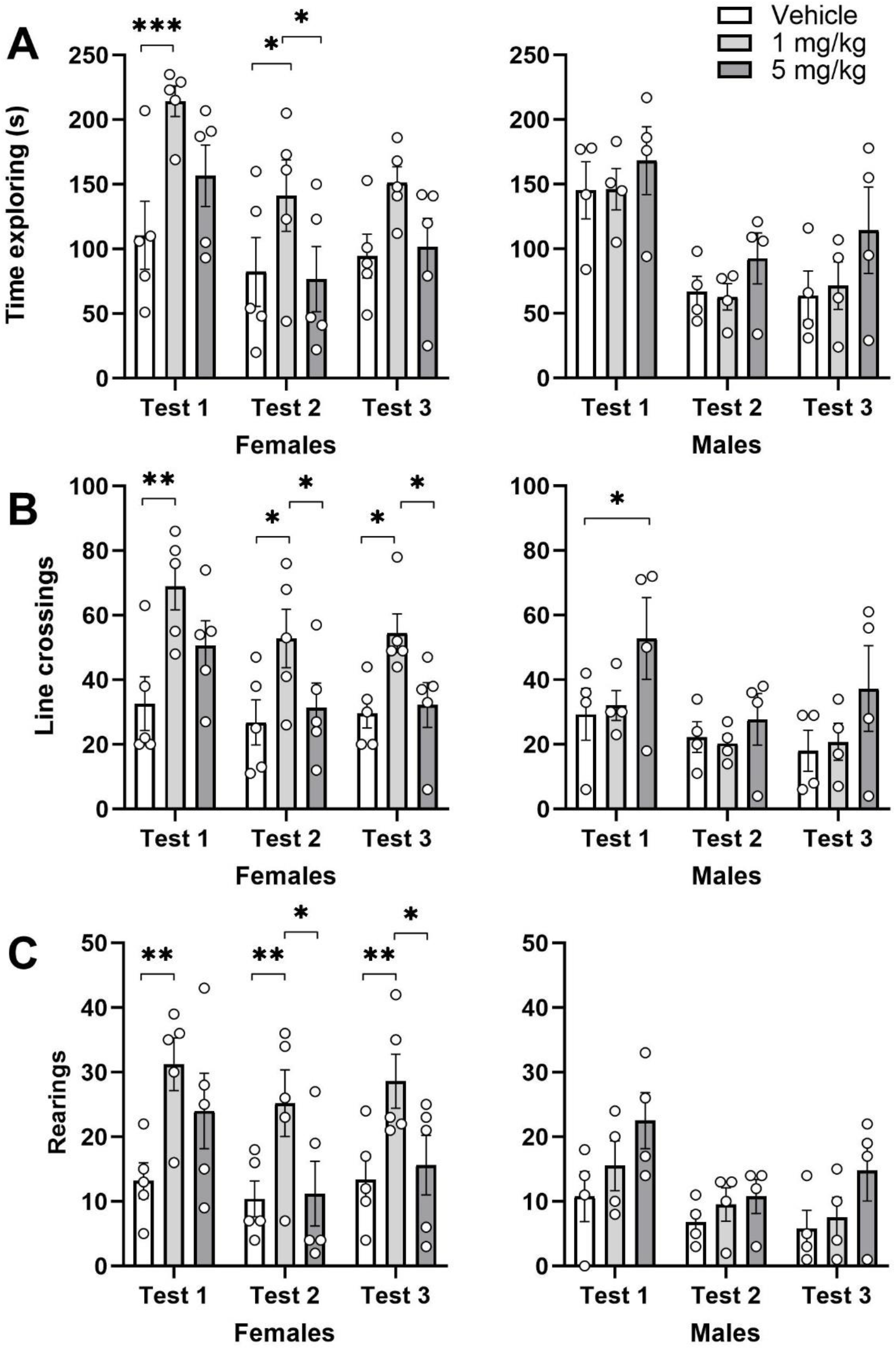
Effect of acute and repeated administration of CHMA1004 on locomotor activity in three observations on an open field. *p<0.05; **p<0.01; ***p<0.001 vs. comparison group.

### Effect of acute administration of CHMA1004 on ultrasonic vocalization during the open field test

Acute CHMA1004 treatment had no statistically significant effect on 50-kHz USVs (Figure 2). Only one rat produced 22-kHz USVs, suggestive that neither testing nor the drug was perceived as anxiogenic. Emission of 50-kHz USVs was also low, with a tendency of more vocalization after CHMA1004 treatment. In conclusion, acute administration of CHMA1004 was without strong negative or positive emotional valence.

### Effect of repeated administration of CHMA1004 on behaviour in elevated zero-maze

Time spent in the open sections of the elevated zero-maze was longer at the second test (Test F1,20=16.7; p<0.001), especially in females. CHMA1004 treatment had no statistically significant effect while the measure tended to be increased (Figure 4).

**Figure 4.**
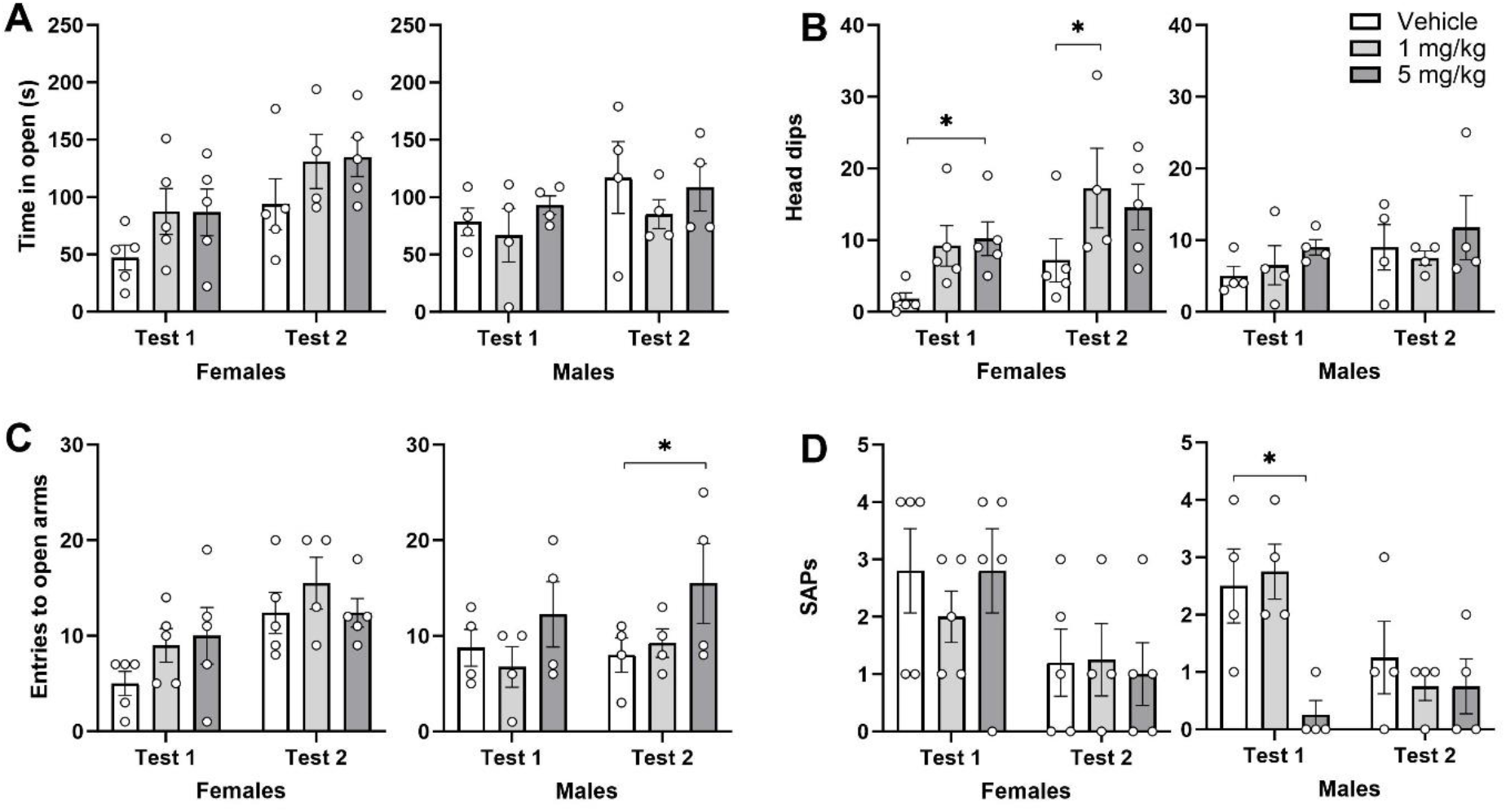
Effect of repeated administration of CHMA1004 in elevated zero-maze test. SAPs, number of stretched-attend postures; *p<0.05 vs. vehicle.

Repeated CHMA1004 treatment increased the number of head-dips at the first zero-maze test (CHMA1004 F2,20=5.35; p<0.05); this effect was more prominent in female rats. The number of entries into the open section was significantly higher at the second testing (Test F1,20=18.6; p<0.001), and was significantly increased in male rats of the CHMA1004 5 mg/kg group as compared to the control at the second elevated zero-maze test (Sex × Test F1,20=5.77; p<0.05). Stretched-attend postures were reduced at the second test (Test F1,20=19.3; p<0.001), however the male CHMA1004 5 mg/kg treatment group had this measure reduced already at the first elevated zero-maze test (Treatment × Sex × Test F2,20=3.09; p=0.067; CHMA1004 F2,20=8.03; p<0.01). In sum, CHMA1004 had an anxiolytic-like effect in after repeated administration.

### Effect of repeated administration of CHMA1004 in social interaction test

In rats housed in groups, social interaction with an animal from another cage was very low (40- 50 s on average) in female rats, and was reduced in the male CHMA1004 5 mg/kg group to the level comparable to females (Sex F1,20=27.4; p<0.001; Treatment × Sex F2,20=3.33; p=0.057; data not shown).

### Effect of repeated administration of CHMA1004 on novelty-induced ultrasonic vocalizations

Not a single rat produced any 22-kHz UVSs during this test, but 50-kHz USVs were emitted by all animals and frequently by most of them. Two-way ANOVA revealed no statistically significant Treatment effect but a signicant Sex effect (F1,21=8.08; p<0.01). Male control rats had higher 50-kHz emission than female controls, but CHMA1004 treatment tended to reduce this to the level of females (Figure 5).

**Figure 5.**
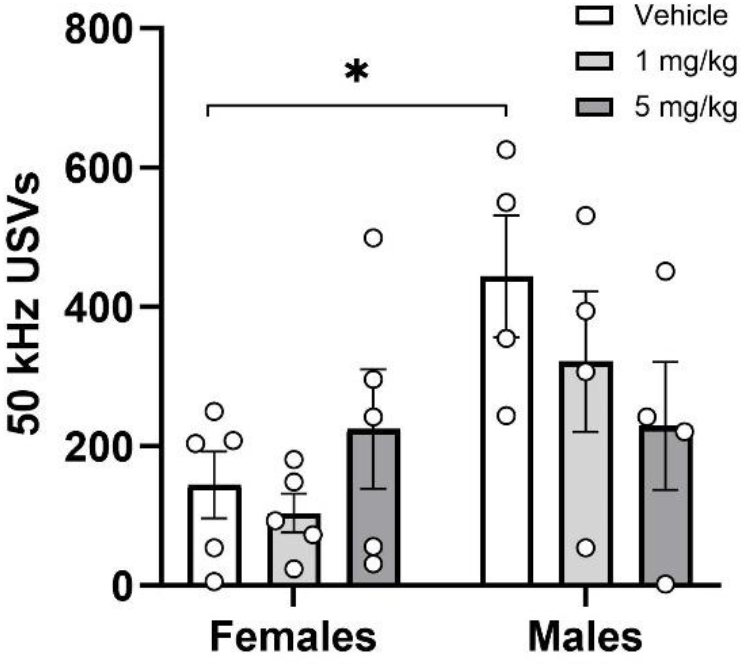
Effect of repeated administration of CHMA1004 on novelty-induced ultrasonic vocalizations.*p<0.05 difference between female and male control groups.

### Effect of repeated administration of CHMA1004 in the forced swimming test

In this screening test for antidepressant potential, CHMA1004 treatment had no significant effect on immobility (Figure 6), but increased swimming (Treatment F 2,21=3.75; p<0.05; Test F1,21=40.6; p<0.001; Sex × Test F1,21=16.5; p<0.001) and reduced struggling (Test F1,21=13.8; p<0.01; Sex × Test F1,21=11.9; p<0.01) in female rats on both days of forced swimming, at both dose levels.

**Figure 6.**
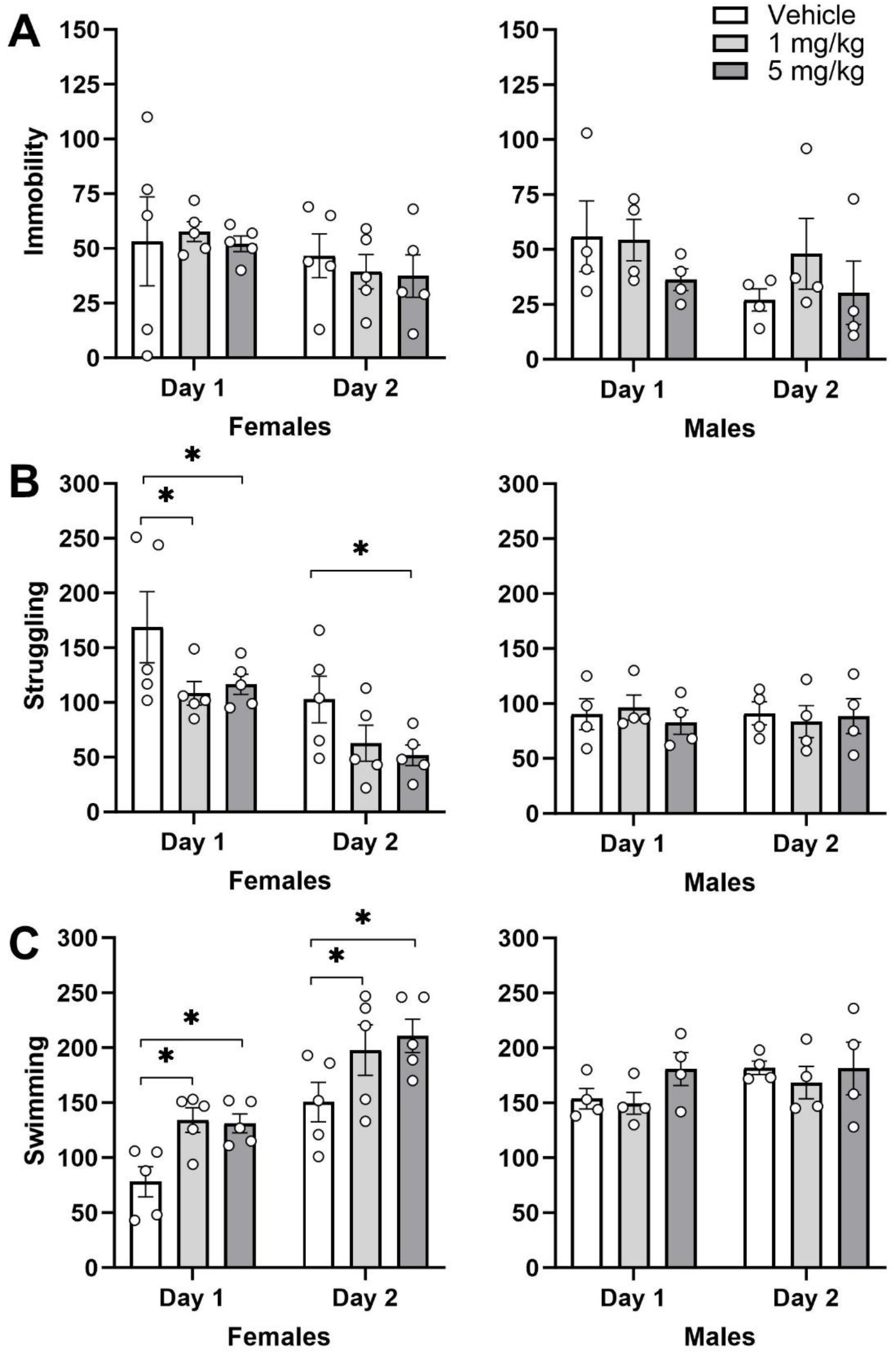
Effect of repeated administration of CHMA1004 in the forced swimming test. *p<0.05 vs. vehicle.

### Effect of repeated administration of CHMA1004 on effects of amphetamine

This was studied in order to assess the potential of RNA m^6^A methylation to cause sensitization to psychostimulant action. After acute administration of amphetamine (1 mg/kg) female rats made more rearings than males (Figure 7A), especially during the latter part of the test (Sex F1,21=27.8; p<0.001; Test F5,105=26.4; p<0.001; Sex × Test F1,21=2.88; p<0.05), for horizontal activity the differences were not statistically significant. In male rats, locomotion tended to be higher in the CHMA1004 (5 mg/kg) group, but this effect was not statistically significant (Figure 7AB).

**Figure 7.**
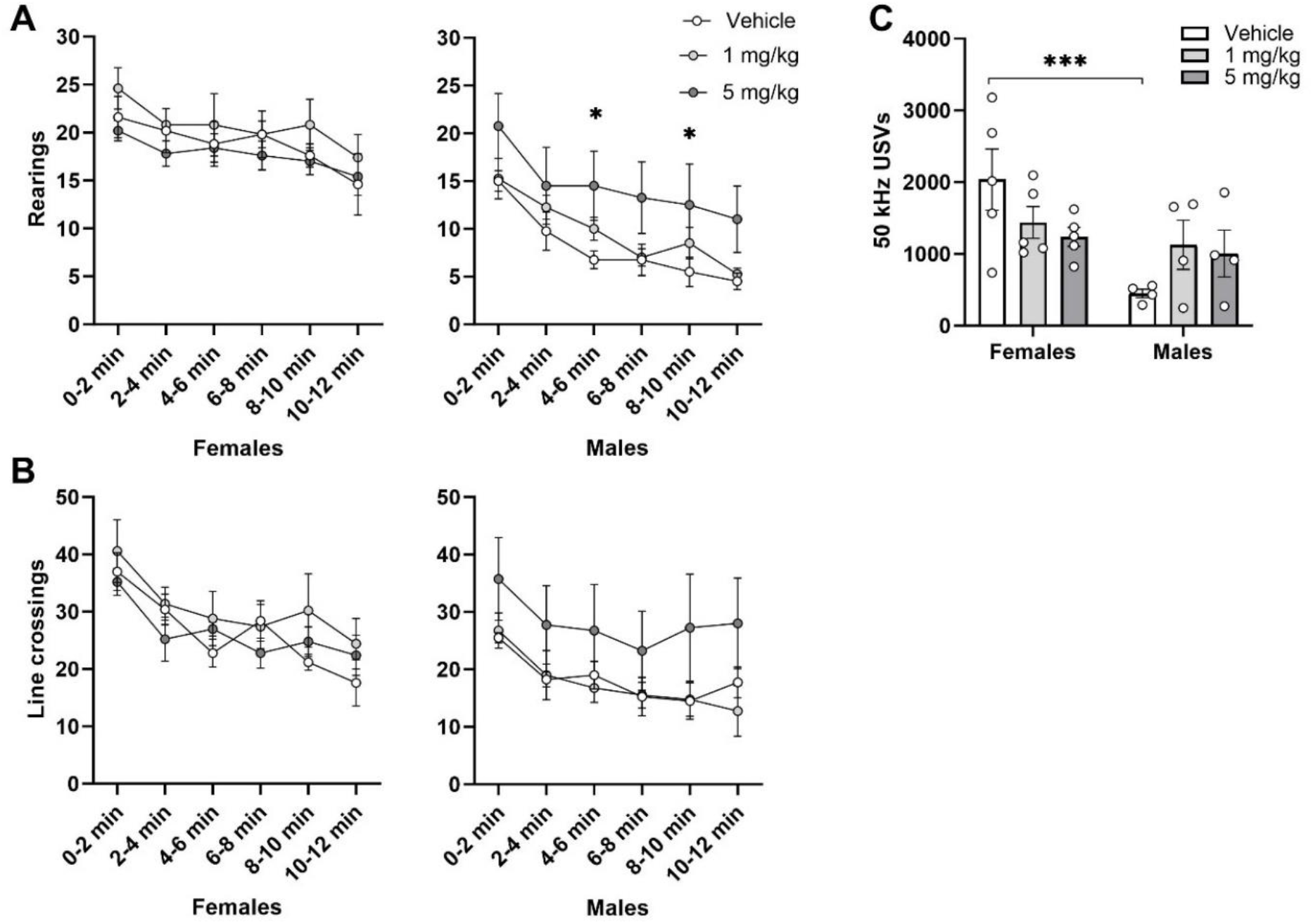
Effects of amphetamine in an open field after repeated administration of CHMA1004. A: number of rearings and B: number of line crossings by 2-min bins; C: total number of 50-kHz ultrasonic vocalizations. *p<0.05 vs. vehicle; ***p<0.001 difference between male and female rats.

After treatment with amphetamine only three male rats and no female rat emitted 22-kHz USVs, but as expected the emission of 50-kHz USVs was robust (Figure 7C). Female rats produced more 50-kHz USVs than males, but this was dependent on CHMA1004 treatment (Sex F1,21=9.17; p<0.01; Treatment × Sex F2,21=3.52; p<0.05). Thus, in male rats subchronic CHMA1004 treatment tended to increase the emission of 50-kHz USVs, but in females this treatment reduced vocalization, so that the large male-female difference in control groups was much smaller in CHMA1004 treatment groups.

### Effect of repeated administration of CHMA1004 on sucrose intake and preference

CHMA1004 treament did not statistically significantly affect sucrose intake or preference (data not shown), but if intake of 11 h was adjusted to body weight, an increase was found in female rats of the CHMA1004, significantly in the 1 mg/kg group, as compared to controls (Figure 8; Sex F1,21=31.2; p<0.001).

**Figure 8.**
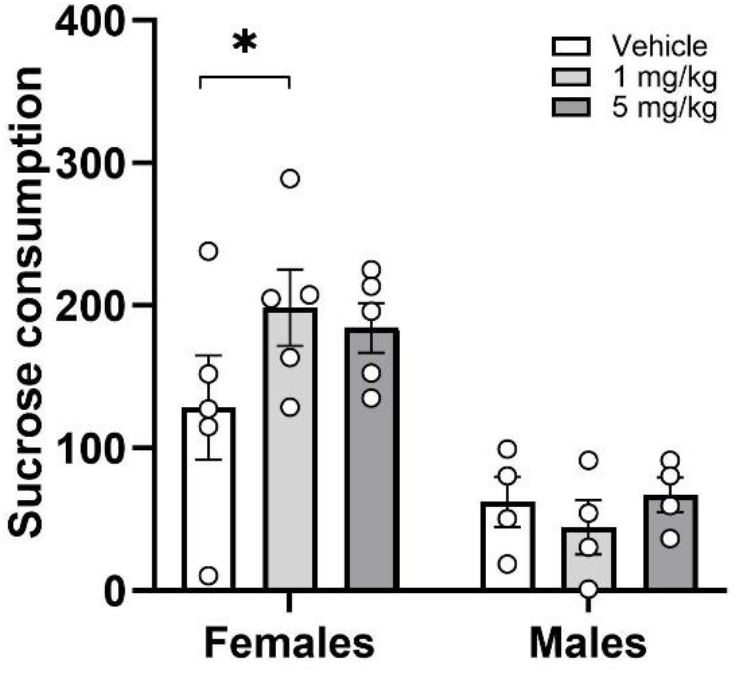
Effect of repeated administration of CHMA1004 on sucrose consumption. 11-h sucrose consumption expressed as g/kg. *p<0.05 vs. vehicle.

### Effect of repeated administration of CHMA1004 on monoamine neurochemistry

Subchronic treatment with most of psychotropic medications causes adaptove changes in one or several monomiane systems, and indeed CHMA1004 affected the levels of monoamine neurotransmitters and their metabolites in several brain regions (Table 1). In striatum, CHMA1004 treatment increased the levels of NMN (Treatment F2,21=4.30; p<0.05; Sex F1,21=3.41; p<0.05) in females and dopamine turnover in males (Treatment × Sex F2,21=4.44; p<0.05). Frontal cortical NMN levels were increased by subchronic treatment with CHMA1004 in males (Treatment F2,21=4.94; p<0.05; Treatment × Sex F2,21=4.73; p<0.05). Similar effect on noradrenaline levels closely missed the conventional level of significance (Treatment F2,21=3.40; p=0.0525), and both NA and NMN tended to be higher also in females at the lower dose of CHMA1004. In locus coeruleus, CHMA1004 treatment reduced HVA levels in female rats, while the effect was opposite in males (Treatment × Sex F(2,21)=4.66; p<0.05). Ventral tegmental area was also dose-dependently affected by CHMA1004 treatment: Levels of noradrenaline and HVA were higher after 1 mg/kg (Treatment F2,21=5.22 and 4.21, respectively; p<0.05; post-hoc tests not significant owing to small groups by sex and large male-female differences), and 5-HIAA levels would be significantly affected in the same manner if one outlier in the vehicle group were excluded (Treatment F2,20=6.42; p<0.01). In hippocampus and in the raphe area, two-way ANOVA did not reveal any statistically significant treatment effect.

**Table 1.** Monoamine levels after two and half weeks of administration of the METTL3/METTL14 activating compound CHMA1004 in male and female rats.

|  | NA | NMN | NMN/NA | DA | DOPAC | DOPAC/DA | 3MT | HVA | 5HT | 5-HIAA | 5-HIAA/5HT |
| --- | --- | --- | --- | --- | --- | --- | --- | --- | --- | --- | --- |
| **Striatum** |  |  |  |  |  |  |  |  |  |  |  |
| *Males* |  |  |  |  |  |  |  |  |  |  |  |
| Vehicle | 4.80 ± 0.74 | 5.79 ± 0.33 | 1.29 ± 0.20 | 75.17 ± 7.47 | 6.34 ± 0.75 | 0.08 ± 0.002 | 4.60 ± 0.59 | 3.69 ± 0.34 | 5.84 ± 0.23 | 2.51 ± 0.10 | 0.43 ± 0.01 |
| CHMA1004<br>1 mg/kg | 4.79 ± 0.64 | 5.11 ± 0.21 | 1.15 ± 0.23 | 73.31 ± 8.26 | 6.66 ± 0.80 | 0.09 ± 0.001 | 4.29 ± 0.30 | 3.78 ± 0.41 | 5.78 ± 0.28 | 2.33 ± 0.13 | 0.40 ± 0.02 |
| CHMA1004<br>5 mg/kg | 6.04 ± 0.92 | 6.19 ± 0.28<sup>#</sup> | 1.09 ± 0.15 | 76.56 ± 7.12 | 7.78 ± 0.47 | 0.10 ± 0.01* | 4.68 ± 0.20 | 4.03 ± 0.34 | 5.86 ± 0.54 | 2.53 ± 0.20 | 0.43 ± 0.03 |
| *Females* |  |  |  |  |  |  |  |  |  |  |  |
| Vehicle | 5.52 ± 0.40 | 5.83 ± 0.33 | 1.07 ± 0.06 | 71.75 ± 5.65 | 7.33 ± 0.76 | 0.10 ± 0.01 | 4.02 ± 0.19 | 4.45 ± 0.23 | 5.89 ± 1.29 | 2.74 ± 0.18 | 0.55 ± 0.10 |
| CHMA1004<br>1 mg/kg | 5.72 ± 0.45 | 6.32 ± 0.38 | 1.12 ± 0.06 | 79.27 ± 7.03 | 6.66 ± 0.74 | 0.08 ± 0.01* | 4.05 ± 0.36 | 4.65 ± 0.54 | 5.60 ± 0.54 | 2.67 ± 0.17 | 0.49 ± 0.03 |
| CHMA1004<br>5 mg/kg | 6.49 ± 0.71 | 7.08 ± 0.40* | 1.13 ± 0.10 | 86.64 ± 6.87 | 7.82 ± 0.90 | 0.09 ± 0.01 | 4.12 ± 0.33 | 5.21 ± 0.42 | 6.61 ± 0.84 | 3.19 ± 0.30 | 0.50 ± 0.04 |
| **Frontal cortex** |  |  |  |  |  |  |  |  |  |  |  |
| *Males* |  |  |  |  |  |  |  |  |  |  |  |
| Vehicle | 6.56 ± 0.63 | 4.46 ± 0.31 | 0.68 ± 0.02 | 0.35 ± 0.05 |  |  | 0.28 ± 0.04 | 0.16 ± 0.03 | 3.07 ± 0.12 | 1.65 ± 0.18 | 0.53 ± 0.04 |
| CHMA1004<br>1 mg/kg | 7.94 ± 0.72 | 6.03 ± 0.44* | 0.76 ± 0.03 | 0.33 ± 0.04 |  |  | 0.34 ± 0.10 | 0.20 ± 0.03 | 3.27 ± 0.11 | 1.50 ± 0.10 | 0.46 ± 0.03 |
| CHMA1004<br>5 mg/kg | 8.04 ± 0.18* | 6.57 ± 0.36** | 0.82 ± 0.05 | 0.37 ± 0.05 |  |  | 0.27 ± 0.04 | 0.20 ± 0.04 | 3.21 ± 0.10 | 1.73 ± 0.16 | 0.54 ± 0.06 |
| *Females* |  |  |  |  |  |  |  |  |  |  |  |
| Vehicle | 7.48 ± 0.11 | 6.10 ± 0.38 | 0.82 ± 0.05 | 0.37 ± 0.03 |  |  | 0.33 ± 0.08 | 0.20 ± 0.04 | 3.75 ± 0.22 | 1.65 ± 0.10 | 0.45 ± 0.04 |
| CHMA1004<br>1 mg/kg | 8.44 ± 0.48 | 6.94 ± 0.45 | 0.84 ± 0.09 | 0.36 ± 0.04 |  |  | 0.42 ± 0.06 | 0.21 ± 0.02 | 3.31 ± 0.19 | 1.54 ± 0.08 | 0.47 ± 0.02 |
| CHMA1004<br>5 mg/kg | 7.50 ± 0.39 | 5.82 ± 0.39<sup>#</sup> | 0.78 ± 0.04 | 0.37 ± 0.04 |  |  | 0.22 ± 0.004 | 0.23 ± 0.02 | 3.53 ± 0.11 | 2.03 ± 0.40 | 0.57 ± 0.11 |
| **Hippocampus** |  |  |  |  |  |  |  |  |  |  |  |
| *Males* |  |  |  |  |  |  |  |  |  |  |  |
| Vehicle | 2.42 ± 0.36 | 0.11 ± 0.02 | 0.05 ± 0.01 | 0.21 ± 0.03 | 0.10 ± 0.01 | 0.48 ± 0.06 |  | 0.05 ± 0.01 | 2.17 ± 0.25 | 1.65 ± 0.18 | 0.77 ± 0.07 |
| CHMA1004<br>1 mg/kg | 2.28 ± 0.12 | 0.08 ± 0.02 | 0.03 ± 0.01 | 0.29 ± 0.09 | 0.10 ± 0.01 | 0.45 ± 0.12 |  | 0.05 ± 0.01 | 1.96 ± 0.13 | 1.43 ± 0.10 | 0.73 ± 0.05 |
| CHMA1004<br>5 mg/kg | 2.18 ± 0.11 | 0.09 ± 0.02 | 0.04 ± 0.01 | 0.18 ± 0.07 | 0.09 ± 0.02 | 0.59 ± 0.10 |  | 0.04 ± 0.01 | 2.36 ± 0.31 | 1.69 ± 0.19 | 0.78 ± 0.17 |
| *Females* |  |  |  |  |  |  |  |  |  |  |  |
| Vehicle | 1.98 ± 0.16 | 0.07 ± 0.01 | 0.04 ± 0.01 | 0.27 ± 0.05 | 0.11 ± 0.01 | 0.44 ± 0.06 |  | 0.06 ± 0.01 | 1.92 ± 0.17 | 1.59 ± 0.06 | 0.86 ± 0.10 |
| CHMA1004<br>1 mg/kg | 2.09 ± 0.18 | 0.07 ± 0.02 | 0.04 ± 0.01 | 0.59 ± 0.19 | 0.18 ± 0.04 | 0.38 ± 0.10 |  | 0.07 ± 0.02 | 1.95 ± 0.14 | 1.69 ± 0.05 | 0.88 ± 0.05 |
| CHMA1004<br>5 mg/kg | 2.20 ± 0.08 | 0.07 ± 0.02 | 0.03 ± 0.01 | 0.28 ± 0.06 | 0.11 ± 0.02 | 0.44 ± 0.07 |  | 0.07 ± 0.02 | 1.87 ± 0.07 | 1.77 ± 0.11 | 0.95 ± 0.06 |
| **Locus coeruleus** |  |  |  |  |  |  |  |  |  |  |  |
| *Males* |  |  |  |  |  |  |  |  |  |  |  |
| Vehicle | 13.26 ±<br>1.45 | 9.00 ± 0.49 | 0.71 ± 0.11 | 0.68 ± 0.03 |  |  |  | 0.37 ± 0.05 | 5.65 ± 0.96 | 4.25 ± 0.86 | 0.76 ± 0.08 |
| CHMA1004<br>1 mg/kg | 17.87 ±<br>2.44 | 10.80 ± 0.72 | 0.63 ± 0.07 | 1.12 ± 0.35 |  |  |  | 1.34 ± 0.06* | 5.68 ± 0.95 | 3.82 ± 0.64 | 0.68 ± 0.03 |
| CHMA1004<br>5 mg/kg | 17.85 ±<br>2.72 | 12.50 ± 3.01 | 0.70 ± 0.13 | 0.94 ± 0.29 |  |  |  | 0.95 ± 0.29 | 6.91 ± 2.17 | 4.08 ± 1.27 | 0.59 ± 0.03 |
| *Females* |  |  |  |  |  |  |  |  |  |  |  |
| Vehicle | 14.53 ± 1.39 | 11.45 ± 1.94 | 0.78 ± 0.11 | 1.20 ± 0.12 |  |  |  | 0.98 ± 0.18 | 9.73 ± 1.65 | 6.38 ± 1.07 | 0.66 ± 0.04 |
| CHMA1004<br>1 mg/kg | 14.79 ± 1.30 | 8.82 ± 1.11 | 0.61 ± 0.10 | 0.79 ± 0.05 |  |  |  | 0.47 ± 0.03* | 5.97 ± 0.53 | 3.55 ± 0.61 | 0.58 ± 0.07 |
| CHMA1004<br>5 mg/kg | 14.26 ± 1.45 | 10.52 ± 0.63 | 0.77 ± 0.09 | 0.78 ± 0.19 |  |  |  | 0.48 ± 0.04* | 7.20 ± 0.61 | 4.30 ± 0.46 | 0.60 ± 0.06 |
| **Raphe** |  |  |  |  |  |  |  |  |  |  |  |
| *Males* |  |  |  |  |  |  |  |  |  |  |  |
| Vehicle | 6.38 ± 0.31 | 0.18 ± 0.06 | 0.03 ± 0.01 | 1.71 ± 0.19 | 0.51 ± 0.06 | 0.30 ± 0.02 |  | 0.62 ± 0.09 | 6.36 ± 0.68 | 7.71 ± 0.59 | 1.27 ± 0.21 |
| CHMA1004<br>1 mg/kg | 5.82 ± 0.52 | 0.12 ± 0.03 | 0.02 ± 0.005 | 1.39 ± 0.09 | 0.38 ± 0.03 | 0.27 ± 0.01 |  | 0.53 ± 0.08 | 6.49 ± 0.22 | 6.58 ± 0.21 | 1.02 ± 0.05 |
| CHMA1004<br>5 mg/kg | 6.22 ± 0.73 | 0.16 ± 0.05 | 0.03 ± 0.01 | 1.43 ± 0.32 | 0.42 ± 0.11 | 0.29 ± 0.02 |  | 0.39 ± 0.10 | 6.01 ± 1.15 | 6.90 ± 0.96 | 1.21 ± 0.14 |
| *Females* |  |  |  |  |  |  |  |  |  |  |  |
| Vehicle | 5.52 ± 1.07 | 0.20 ± 0.05 | 0.04 ± 0.01 | 1.74 ± 0.42 | 0.45 ± 0.08 | 0.28 ± 0.04 |  | 0.45 ± 0.10 | 6.62 ± 0.89 | 6.34 ± 0.81 | 0.96 ± 0.05 |
| CHMA1004<br>1 mg/kg | 6.09 ± 0.15 | 0.15 ± 0.03 | 0.03 ± 0.005 | 1.48 ± 0.19 | 0.40 ± 0.04 | 0.27 ± 0.02 |  | 0.47 ± 0.05 | 7.16 ± 0.25 | 7.04 ± 0.77 | 0.98 ± 0.08 |
| CHMA1004<br>5 mg/kg | 6.11 ± 0.23 | 0.14 ± 0.03 | 0.02 ± 0.01 | 1.25 ± 0.07 | 0.41 ± 0.07 | 0.33 ± 0.05 |  | 0.42 ± 0.04 | 7.32 ± 0.31 | 7.64 ± 0.30 | 1.05 ± 0.05 |
| **Ventral tegmental area** |  |  |  |  |  |  |  |  |  |  |  |
| *Males* |  |  |  |  |  |  |  |  |  |  |  |
| Vehicle | 4.50 ± 0.86 | 0.12 ± 0.02 | 0.03 ± 0.005 | 2.55 ± 1.22 | 0.71 ± 0.36 | 0.31 ± 0.08 | 0.42 ± 0.15 | 0.82 ± 0.27 | 6.66 ± 1.08 | 3.93 ± 0.93 | 0.61 ± 0.11 |
| CHMA1004<br>1 mg/kg | 8.70 ± 1.89<sup>a</sup> | 0.35 ± 0.13 | 0.03 ± 0.003 | 4.51 ± 1.48 | 1.06 ± 0.29 | 0.26 ± 0.03 | 1.07 ± 0.93 | 1.53 ± 0.32 | 7.14 ± 1.29 | 5.95 ± 0.66 | 0.69 ± 0.14 |
| CHMA1004<br>5 mg/kg | 6.26 ± 2.18 | 0.34 ± 0.24 | 0.03 ± 0.007 | 1.78 ± 0.50 | 0.85 ± 0.29 | 0.63 ± 0.28 | 0.48 ± 0.32 | 0.93 ± 0.42 | 8.09 ± 1.58 | 3.80 ± 0.11 | 0.60 ± 0.14 |
| *Females* |  |  |  |  |  |  |  |  |  |  |  |
| Vehicle | 5.28 ± 0.41 | 0.06 ± 0.01 | 0.60 ± 0.42 | 3.71 ± 1.43 | 0.71 ± 0.27 | 0.19 ± 0.01 | 0.39 ± 0.13 | 0.83 ± 0.15 | 6.81 ± 1.28 | 6.10 ± 1.60 | 1.12 ± 0.49 |
| CHMA1004<br>1 mg/kg | 8.02 ± 1.54 | 0.21 ± 0.12 | 0.89 ± 0.86 | 6.16 ± 1.38 | 1.45 ± 0.36 | 0.23 ± 0.01 | 1.03 ± 0.53 | 1.65 ± 0.39 | 10.62 ± 2.26 | 6.21 ± 0.84 | 0.64 ± 0.07 |
| CHMA1004<br>5 mg/kg | 4.79 ± 0.61 | 0.08 ± 0.02 | 0.74 ± 0.45 | 3.75 ± 1.15 | 1.04 ± 0.26 | 0.38 ± 0.17 | 0.67 ± 0.46 | 0.79 ± 0.17<sup>#</sup> | 6.38 ± 1.36 | 4.08 ± 0.50 | 0.69 ± 0.08 |

Monoamine levels are expressed as pmol/mg tissue. \* $p < 0.05$ ; \*\* $p < 0.01$ vs. vehicle; <sup>#</sup> $p < 0.05$ vs. C4 (1 mg/kg); <sup>‡</sup> $p = 0.055$ vs. vehicle.

### Extracellular levels of dopamine in striatum

To assess whether METTL3/14 activation could be triggering dopamine release we conducted a separate microdialysis experiment. As expected, acute administration of saline had no effect of striatal dopamine levels, while amphetamine (0.5 mg/kg) increased dopamine release in male (F51,187=6.88; p<0.001) and female (F34,170=5.58; p<0.001) rats. CHMA1004 at either dose did not elicit any significant effect on extracellular dopamine levels in striatum (Figure 9).

**Figure 9.**
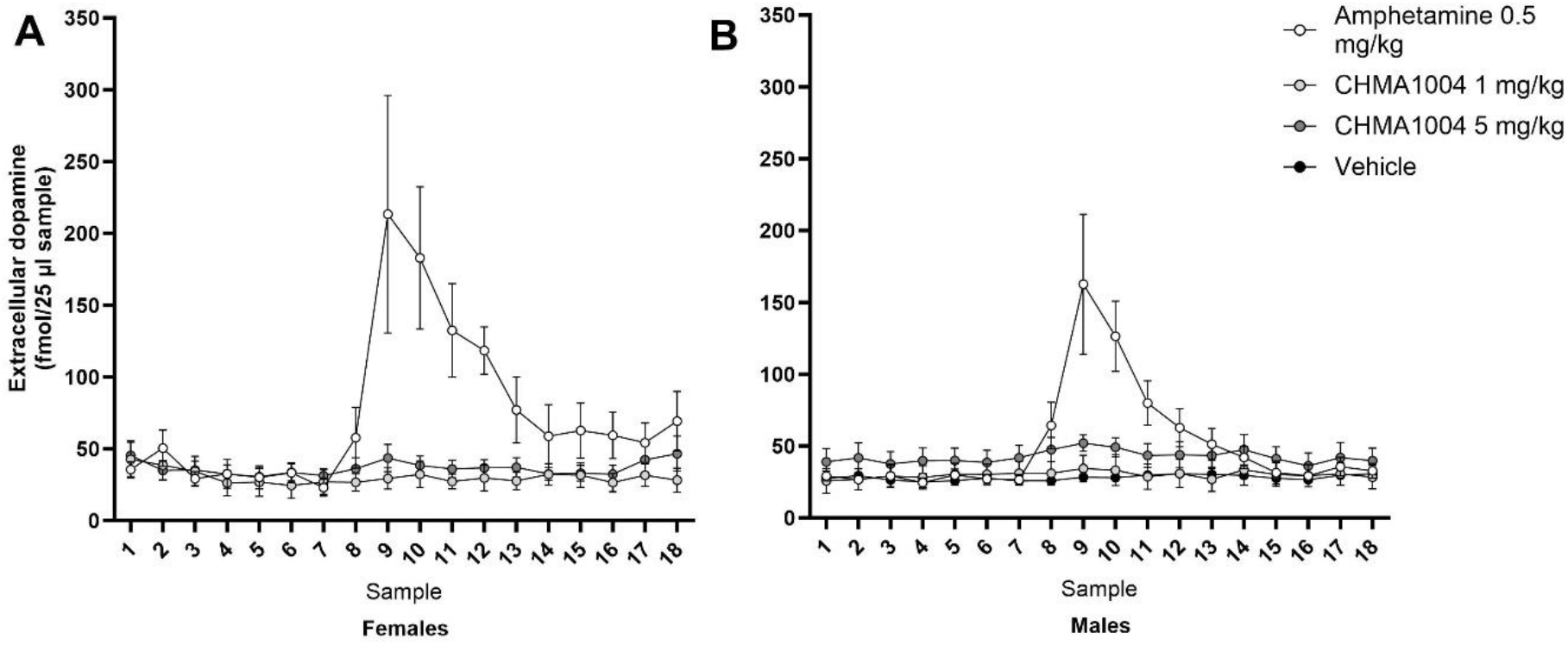
Extracellular levels of dopamine after acute administration of amphetamine or CHMA1004. A: Female rats; B: Male rats.

### Effect of subchronic administration of CHMA1004 on striatal gene expression

Because it is not known which epitranscriptomic changes should be expected to be most responsive to METTL3/METTL14 activation, a hypothesis free approach, RNA sequencing was performed. Differential expression analysis resulted in statistically significant change (Wald test p<0.05) of 172 and 120 genes following daily 1 mg/kg and 5 mg/kg of CHMA1004, respectively (Supplementary Table 1). Of these genes, seven were overlapping (Galns, Hps3, Misp3, Nbl1, Samd4a, Slc28a2, Spon2), with similar change in expression. Among the genes that were affected with the highest statistical significance (Figure 10), most were of specific interest, e.g., Acsm5, Peli1, Pcdhb5, Psmb9, Fgr, Lrp10, Grb83, Hes1, Vwa5b2, Mx2, Serpinf2, Nbl1 and Mdga1; of other differentially expressed genes, Nxph3 would deserve particular attention (see Discussion).

**Figure 10.**
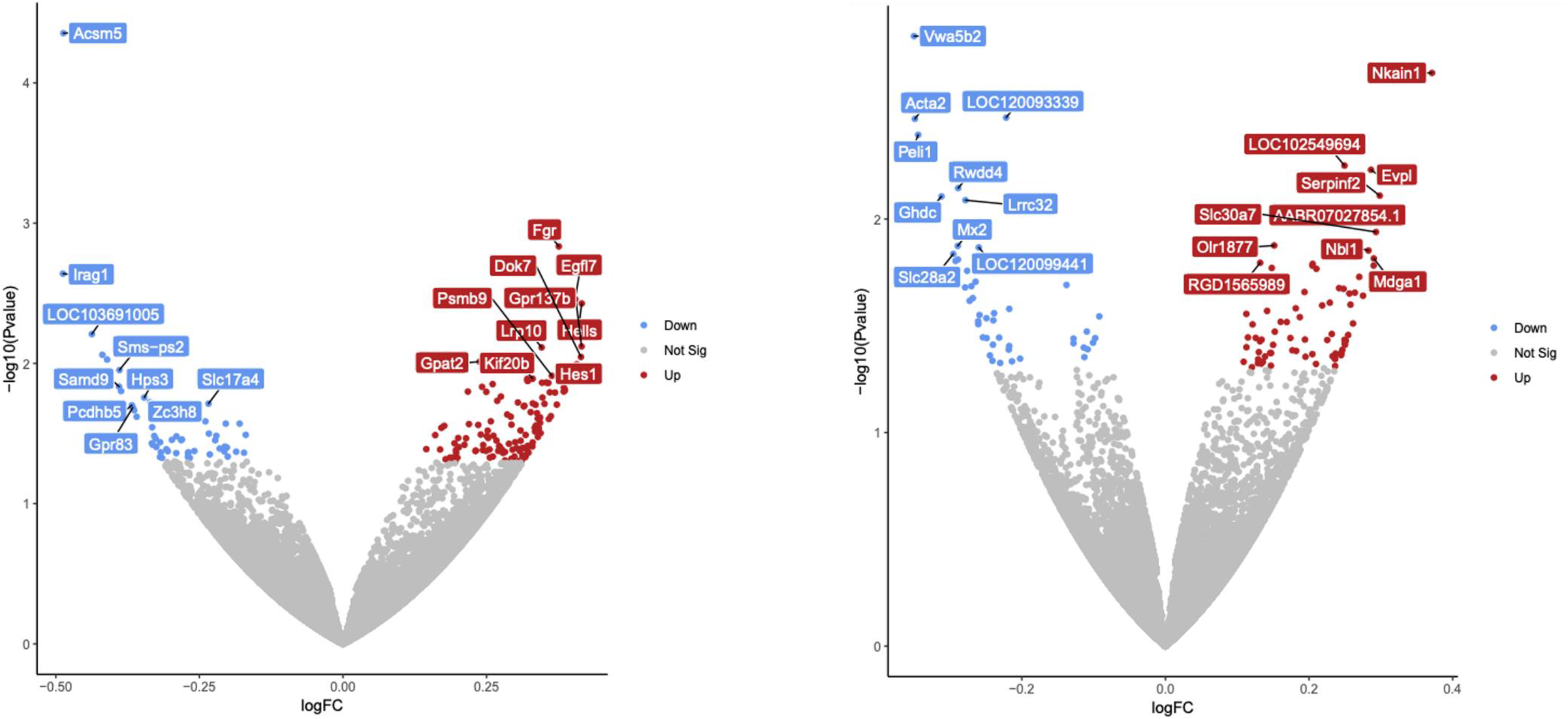
Effect of repeated administration of CHMA1004 on striatal gene expression. Volcano plots of the differentially expressed gene analysis with DESeq2 between control and CHMA1004 1 mg/kg (left) or 5 mg/kg (right) treated female rats. Significantly differentially expressed genes (p<0.05) are labelled blue (downregulated genes) or red (upregulated genes). The names of the top 10 genes with the lowest p-values are shown.

Pathway analyses (Supplementary Table 2) suggested the involvement of e.g., phospholipase C-activating G-protein coupled receptor signalling, positive regulation of phosphatidylinositol 3-kinase signalling metabolic pathways, cell adhesion, and mitochondrial matrix in the action of CHMA1004.

## Discussion

This study revealed that acute and subchronic administration of a novel METTL3/METTL14 activator compound CHMA1004 can elicit activating and anxiolytic-like effects in the rat. Subchronic treatment with CHMA1004 affected monoaminergic neurotransmission in the brainstem and forebrain areas. These effects were region- and dose-dependent and not all identical in male and female animals, but in general were suggestive of an increase in catecholamine function. Subchronic CHMA1004 treatment also elicited many alterations in striatal gene expression that are relevant to stress response.

Acute treatment effect of CHMA1004 was assessed in an open field test. Increased locomotor activity that was observed can be indicative of anxiolytic action, but also of a psychostimulant effect. Therefore, we simultaneously recorded ultrasonic vocalizations, as classic psychostimulants typically elicit 50-kHz USVs (Simola et al., 2010; Kõiv et al., 2019; Wöhr, 2022). No strong psychostimulant effect was observed, suggesting that the increased locomotor activity was reflective either reduced anxiety or a non-dopaminergic psychostimulant effect. On the other hand, rats did not produce 22-kHz USVs. Previously, administration of neuropeptide S that also displays mixed anxiolytic/psychostimulant profile in exploration-based tests (Xu et al., 2004; Rizzi et al., 2008) was found to elicit substantial 22-kHz vocalization revealing predominantly negative emotional response behind behavioural activation (Kõiv et al., 2021). CHMA1004 did, in contrast, display an anxiolytic-like and stimulatory effect without any significant affective response in terms of USV production. That the effect of acute administration of CHMA1004 was anxiolytic, was further suggested by the reduction of defecations on open field in male rats. Previously, acute restraint stress was shown to rapidly downregulate Mettl3 expression in the prefrontal cortex and amygdala of mice (Engel et al., 2018). That the METTL3/METTL14 activating compound CHMA1004 elicited an anxiolytic effect in conditions of novelty stress suggests that RNA m^6^A methylation is involved in active coping with stress.

Repeated treatment with CHMA1004 did not appear to produce significant tolerance to its anxiolytic-like/activating effect. The anxiolytic effect of CHMA1004 was further substantiated by results of the elevated zero-maze test where stretched-attend postures were reduced, open quadrant entries and head-dips increased. At the first glance the tendency of decrease in social interaction time in male rats may appear as contradictory to an anxiolytic effect, but the test was not conducted as validated for this purpose: We could not use single housing in this experiment because this would sensitize animals to stressful stimulation. It is likely that at this locomotor activating dose CHMA1004 increased exploration of the novel physical environment on account of social behaviour. Similar is the effect of amphetamine in the social interaction test (File and Hyde, 1979; unpublished results from this laboratory).

The forced swimming test is the preferred screening tool for novel drugs with antidepressant potential. Subchronic CHMA1004 treatment did not affect immobility but increased swimming and reduced struggling in female rats. The characteristic response to the first generation antidepressant drugs such as tricyclics is reduction of immobility (Porsolt et al., 1978) while the selective serotonin reuptake inhibitors increase swimming (Armario et al., 1988). It should be noted that in the forced swimming test used as a screening tool mostly acute drug treatment after the first swimming session is applied. In the present study the drug had been repeatedly administered before the test and immobility was relatively low in these repeatedly handled rats. As expected, swimming increased and struggling decreased when CHMA1004 treated animals animals were submitted to forced swimming the second time. Previous studies applying chronic stress or partial serotonergic denervation have suggested that a behavioural activation that is present already at the first swimming session is indicative of higher expression of impulsivity (Harro et al., 2001). Thus, the present results could be interpreted as subchronic CHMA1004 treatment presenting an impulsivity-reducing effect. This possibility should be examined further in behavioural tests specifically targeting different aspects of impulsivity (Dalley and Robbins, 2017). In another test of antidepressant prediction, the sucrose preference test, sucrose preference was not affected but consumption as adjusted to body weight was increased by CHMA1004 treatment in females. Again, the test is validated in conditions of chronic stress induced reduction of sucrose intake, and as part of the present test battery informs little about potential antidepressant effect but suggests that subchronic CHMA1004 treatment is not stressful/anhedonic and rather may increase the appetitive value of carbohydrates. In this context it is also relevant that long-term treatment with CHMA1004 did not increase adrenal weight, again suggesting that this treatment was not strongly stressful; indeed, it may be important to notice that at the doses that were behaviourally most effective (1 mg/kg in females and 5 mg/kg in males, respectively), a tendency of reduction of adrenal weight appeared.

Fifty-kHz USVs are indicative of positive affect in the rat (Burgdorf et al., 2001; Brudzynski and Granon 2023) and in adult animals the most powerful stimulus to elicit this response is psychostimulant drugs such as amphetamine. Novel home-cage like environment can also elicit the 50-kHz response. Subchronic CHMA1004 treatment did not modify the effect of either novel cage or amphetamine treatment significantly, but statistical power to detect its effect may have been too low, because substantial differences between male and female control groups appeared. A general limitation in this study is the number of animals of each sex was small. This was to reduce the overall number of rats used, and simultaneously provide comparison of male and female animals. It was not foreseen that dose-dependency varies between males and females, and the significantly larger body weight of males may have been a contributory factor to this. Nevertheless, it can be concluded that CHMA1004 did not elicit substantial psychostimulant-like effects or produced sensitization on its own. CHMA1004 treatment enhanced locomotor activity in response to amphetamine challenge in male rats, but their activity remained below of females. Altogether the potential of CHMA1004 to increase amphetamine-like stimulant responses appeared low.

Recent studies have highlighted the fact that dysregulation of m^6^A mRNA methylation occurs in neurodegeneration, and dopamine neurons are dependent on proper regulation of m^6^A mRNA methylation (Chen et al., 2019). Inhibition of m^6^A RNA demethylases Fto and Alkbh5, which results in increased m6A levels, can support the survival of dopamine neurons (Selberg et al., 2021). Furthermore, the METTL3/METTL14 activating compound CHMA1004 was also found to support survival of cultured dopamine neurons that were deprived of growth factors and to protect neurons from 6-OHDA toxicity (Yu et al., 2023). Importantly, intrastriatal administration of CHMA1004 improved motor behaviour and increased tyrosine hydroxylase positive fibre density in mice after 6-OHDA lesion; it also protected human cultured dopamine neurons in that study. Hence, we measured monoamine levels *ex vivo* in striatum after long-term administration of CHMA1004, and having observed alterations in catecholamines, performed further measurements in a few other brain regions. The most prominent effect of subchronic treatment with CHMA1004 was observed in VTA at the lower dose level. Dopaminergic measures tended to be altered also in the other brainstem regions and several alterations were observed in measures of catecholaminergic neurotransmission, mostly increasing with dose. These results suggest a complex reorganization of monoamine systems by a METTL3/METTL14 activating drug, possibly primarily in the dopamine neurons. Acute CHMA1004 treatment did however not produce any increase in striatal extracellular dopamine levels, as might have been suggested by its behaviourally stimulatory effects, which is consistent with no increase in 50-kHz ultrasonic vocalizations after either acute or subchronic CHMA1004 treatment.

Most of proposed pathogenetic mechanisms of depression, relating to neurogenesis, activity of the HPA axis, glial function, neuroinflammation and synaptic transmission have all been associated with mRNA m^6^A methylation (Li C et al., 2023). One genetic association study could link a variance in the RNA m^6^A demethylase encoding gene *ALKBH5* with major depression in a large sample (Du et al., 2015) and another associated with depression an *FTO* variant (Samaan et al., 2013). The biochemical pathways involved in the association of mRNA m^6^A methylation with anxiety or depression-related behavioural changes have however remained completely unknown. To extend knowledge beyond the monoamine systems and to analyse what mRNAs are regulated by METTL3/METTL14 activator CHMA1004 we also performed RNA sequencing in striatum in female rats.The low and high dose of CHMA1004 resulted in a different landscape of differential gene expression, suggesting that response to subchronic drug treatment involved dose-dependent adaptive changes that had also been apparent in behavioural and biochemical effects; however, expression of several genes was altered in a similar manner. Many effects of CHMA1004 on gene expression agree with what is known of m^6^A in neurodegeneration and affective disorders (see Introduction), and the capacity of the drug to protect dopamine neurons (Yu et al., 2023) as well as its behavioural effects in the present study. For example, Galns and Samd4a have been implicated in neuronal viability and plasticity; Galns that encodes a lysosomal exohydrolase has been considered a gene related to Parkinson’s disease (Wettergren et al., 2012) and Samd4a is a RNA binding protein that can bind to a variety of target mRNAs through stem-loop structures, also known as Smaug recognition elements (SREs). Notably, Samd4a can regulate the mRNA stability, degradation and translation (Wang and Zhang, 2023). Fgr (Gardner-Rasheed feline sarcoma viral (v-fgr) oncogene homolog) and Psmb9 (proteasome subunit beta type-9), two innate immunity genes, both of which were affected by CHMA1004 treatment, each were recently associated with specific genetic forms of Parkinson’s disease (Shani et al., 2023). Another differentially expressed gene, Lrp10 (low-density lipoprotein receptor-related protein 10), has similarly been linked to Parkinson’s disease but also other neurodegenerative diseases, whereas its role has been attributed to astrocytic vesicle trafficking (Grochowska et al., 2021), and its contribution to neurodegeneration suggested to be significant specifically in females (Guo et al., 2023). CHMA1004 treatment elicited upregulation of several cognition-related genes such as Acta2, Mx2, Spon2 and Peli1. Peli1, encoding Pellino E3 ubiquitin protein ligase 1, was recently shown to mediate methamphetamine-induced neuroinflammation whereas its downregulation was protective (Xu et al., 2023); similarly, the reduction of Peli1 function reduced the lipopolysaccharide-induced inflammatory response in microglia (Dai et al., 2019). Given that neuroinflammatory mechanisms are set in the brain by stress (Black, 2002), the reduction of Peli1 expression by CHMA1004 treatment may reflect a drug effect facilitating adaptive stress response to the experimental procedures. Mx2, an interferon-inducible GTP-binding protein, has been associated with the beneficial effect of physical exercise in a mouse model of Alzheimer’s disease (Widjaya et al., 2023), and Spon2 that can induce neurite outgrowth with learning effects in monkeys (Matsunaga et al., 2015). Serpinf2 that encodes a serine protease inhibitor has a link to Alzheimer’s disease (De Jager et al., 2014), but was recently found as differentially expressed in a rat model of depression (Liao et al., 2023). Depression-related is also Mdga1, encoding a cell adhesion molecule (Li et al., 2022), as well as a few other differentially expressed genes. Pcdhb5 or protocadherin beta-5 has been reported as a gene with altered expression in suicide victims (Thalmeier et al., 2008). It belongs to the group of cadherin-like cell adhesion molecules and knowledge on its function is limited but likely to include the establishment of specific neural connections. Nbl1, a candidate neuroblastoma suppressor gene has also been associated with suicide (Han et al., 2023).

In turn, regulation of Vwa5b2 at the DNA methylation level has been related to lithium response in bipolar disorder (Marie-Claire et al., 2020). Yet another prominent effect of CHMA1004 was downregulation of Acsm5 that encodes a mitochondrial enzyme that catalyzes the activation of fatty acids by coenzyme A. While very little is known of the brain function of Acsm5, it has been identified as a top attention-deficit/hyperactivity disorder (ADHD)-related gene in a bioinformation transcriptome study (Qi et al., 2019), and ADHD is a vulnerability factor for several other psychiatric disorders including anxiety disorders, depression and bipolar disorder (Chen Q et al., 2018). A few other genes with substantially altered expression by CHMA1004 treatment have been linked to anxiety and stress response: expression of Hes1, an essential Notch signalling effector was associated with neuronal excitability in amygdala and with anxiety in the mouse (Matsuzaki et al., 2019), and knockdown of Gpr83, a glucocorticoid-regulated receptor has been associated with anxiety in female mice and a decrease in receptor expression to dexamethasone treatment, whereas anxiety was reduced in males (Fakira et al., 2021). Indeed, a thorough study on the role of m^6^A methylation in stress response regulation reported that the response of m^6^A levels upon glucocorticoid receptor stimulation was altered in patients with major depression (Engel et al., 2018). In that study, one of the most prominent alterations by knocking out the gene responsible for m^6^A methylation, Mettl3, was downregulation of Nxph3, encoding neurexophilin 3. The METTL3/METTL14 activator CHMA1004, in contrast, increased the expression of Nxph3 in the present study. Neurexophilin 3 is a ligand of synaptic α-neurexins thus contributing to the regulation of neurotransmitter release, and it has a role in sensorimotor gating and motor coordination (Beglopoulos et al., 2005).

The anxiolytic-like effects of a METTL3/METTL14 activating compound appear as inconsistent with findings that downregulation of Fto and increased m^6^A methylation can lead to anxiety- and depression-like behaviours in mice (Liu et al., 2021; Wang XL et al., 2022). In these studies, the FTO-related anxious behaviours were associated with mRNA methylation and expression, respectively, of ß2-adrenoceptor in hippocampus or matrix metalloprotease 9 (MMP-9) in anterior cingulate cortex (ACC). In our study neither Adrb2 nor Mmp9 expression was altered by CHMA1004 in striatum. While we did not examine the same brain regions, the contrasting behavioural findings suggest that METTL3/METTL14 activation by systemic administration of a small molecule compound affects RNA m^6^A methylation dissimilarly from downregulation of Fto by other type of interventions. Another study demonstrated that increased RNA m^6^A methylation of fatty acid amide hydrolase mRNA by reducing the function of ALKBH5 attenuated depressive-like behaviours (Huang et al., 2020), which is in line with our findings and further suggests that some of the multiple mechanisms recruited in modifying RNA m^6^A methylation have a potential in the management of psychiatric disorders. Furthermore, treadmill exercise was shown to counteract the anxiety-provoking effects of chronic stress in an RNA m^6^A methylation dependent manner (Yan et al., 2023), providing support to the notion that METTL3/METTL14 activating drugs have anxiolytic and activating effects by promoting physiological mechanisms.

Conclusively, the first-in-class METTL3/METTL14 activator compound methyl-piperazine-2- carboxylate (CHMA1004) elicited anxiolytic-like effects after acute systemic administration, and its anxiolytic-like and activating properties were consistently observed in a battery of behavioural tests during subchronic treatment. While behaviourally activating, CHMA1004 did not exhibit strong psychostimulant-like properties. Subchronic administration of CHMA1004 increased locomotor activity and led to alterations in catecholaminergic neurotransmission and changes in striatal gene expression linked to dopamine neuron viability, neurodegeneration, depression, anxiety and stress response. Thus, pharmacological activation of RNA m^6^A methylation has potential for neuropsychiatric drug development.

## Acknowledgements

This study was funded by the Estonian Research Council (PRG1213) and the TLU Centre of Excellence in Behavioural and Neural Sciences and the School of Natural Sciences and Health Research Fund. JH was supported by Visiting Professorship of the Sigrid Jusélius Foundation. TO was supported by the European Union through Horizon 2020 research and innovation programme under grant number 810645. IT was funded by EU Marie Sklodowska Curie postdoctoral fellowship 101068830. MK and MS were supported from the Cure Parkinson’s Trust, UK grant (project code MS02).

