## Supplementary Table 1 for "Effect of RNA m^6^A methyltransferase activation by a low molecular weight compound on anxiety- and depression-related behaviours, monoamine neurochemistry and striatal gene expression in the rat"

**Supplementary Table 1.** List of differentially expressed genes (p<0.05) based on RNA-seq analysis.

**CHMA1004 1 mg/kg**

| **Gene symbol** | **Average norm read counts Ctrl** | **Average norm read counts C4 (1mg/kg)** | **P-value** | **Fold change** |
| --- | --- | --- | --- | --- |
| *Klhl10* | 0.88 | 23.50 | 0.04 | 26.61 |
| *Amn* | 0.67 | 13.50 | 0.03 | 20.13 |
| *LOC102549654* | 0.92 | 14.62 | 0.03 | 15.81 |
| *Ankar* | 1.11 | 15.18 | 0.03 | 13.68 |
| *Gpat2* | 1.56 | 19.26 | 0.01 | 12.34 |
| *Fat2* | 3.08 | 30.79 | 0.02 | 9.99 |
| *ENSRNOG00000065049* | 1.16 | 9.22 | 0.04 | 7.97 |
| *Fgr* | 5.42 | 36.36 | 0.00 | 6.71 |
| *Baiap2l1* | 4.21 | 27.92 | 0.01 | 6.63 |
| *Myh8* | 4.70 | 25.88 | 0.02 | 5.51 |
| *Polq* | 3.46 | 18.94 | 0.04 | 5.48 |
| *Snora52* | 2.71 | 14.75 | 0.03 | 5.44 |
| *Mirlet7b* | 4.92 | 26.08 | 0.01 | 5.30 |
| *LOC102548186* | 9.34 | 49.43 | 0.05 | 5.29 |
| *ENSRNOG00000069955* | 6.23 | 32.24 | 0.03 | 5.17 |
| *ENSRNOG00000069103* | 2.09 | 9.99 | 0.05 | 4.79 |
| *ENSRNOG00000069541* | 5.40 | 25.46 | 0.04 | 4.71 |
| *ENSRNOG00000065542* | 4.33 | 19.86 | 0.04 | 4.58 |
| *Nmur2* | 3.87 | 17.39 | 0.05 | 4.49 |
| *AABR07022168.1* | 3.91 | 17.50 | 0.04 | 4.47 |
| *Slc44a3* | 6.63 | 29.63 | 0.03 | 4.47 |
| *LOC100362611* | 4.76 | 20.18 | 0.05 | 4.24 |
| *Bard1* | 2.50 | 10.49 | 0.04 | 4.20 |
| *Treml1* | 5.94 | 24.80 | 0.03 | 4.18 |
| *Asic3* | 5.54 | 22.60 | 0.05 | 4.08 |
| *Mrgprf* | 4.70 | 18.94 | 0.05 | 4.03 |
| *Pmfbp1* | 5.86 | 22.95 | 0.02 | 3.91 |
| *ENSRNOG00000067429* | 5.69 | 20.27 | 0.03 | 3.56 |
| *Ttc22* | 18.90 | 66.49 | 0.03 | 3.52 |
| *Erich5* | 8.36 | 27.90 | 0.03 | 3.34 |
| *Asgr1* | 5.69 | 18.34 | 0.04 | 3.22 |
| *Cdk1* | 14.79 | 47.66 | 0.04 | 3.22 |
| *Pole2* | 7.50 | 24.04 | 0.03 | 3.20 |
| *Rdh16* | 14.16 | 44.43 | 0.01 | 3.14 |
| *Chrne* | 8.37 | 26.12 | 0.01 | 3.12 |
| *ENSRNOG00000069802* | 6.54 | 20.17 | 0.03 | 3.08 |
| *ENSRNOG00000071122* | 14.43 | 44.12 | 0.04 | 3.06 |
| *ENSRNOG00000062478* | 18.08 | 54.19 | 0.02 | 3.00 |
| *Kif20b* | 15.90 | 47.53 | 0.01 | 2.99 |
| *Hells* | 28.49 | 84.28 | 0.00 | 2.96 |
| *Cpg1* | 7.74 | 22.77 | 0.03 | 2.94 |
| *Misp3* | 6.76 | 19.77 | 0.02 | 2.92 |
| *Snord104* | 7.61 | 22.22 | 0.04 | 2.92 |
| *Foxs1* | 17.71 | 49.80 | 0.02 | 2.81 |
| *Zfp101* | 11.81 | 31.71 | 0.04 | 2.69 |
| *Mtbp* | 15.81 | 41.87 | 0.02 | 2.65 |
| *Tekt4* | 13.19 | 34.93 | 0.02 | 2.65 |
| *Il2rg* | 13.34 | 32.77 | 0.04 | 2.46 |
| *Psmb9* | 35.99 | 88.29 | 0.01 | 2.45 |
| *RGD1561440* | 29.96 | 73.21 | 0.02 | 2.44 |
| *Oprd1* | 28.47 | 68.70 | 0.05 | 2.41 |
| *Riiad1* | 27.78 | 66.69 | 0.04 | 2.40 |
| *Fam178b* | 14.93 | 35.22 | 0.02 | 2.36 |
| *Fign* | 24.25 | 56.70 | 0.04 | 2.34 |
| *A3galt2* | 30.68 | 70.71 | 0.02 | 2.30 |
| *Ltb* | 30.83 | 70.11 | 0.04 | 2.27 |
| *Spata1* | 25.17 | 56.69 | 0.03 | 2.25 |
| *AC135409.1* | 67.09 | 150.13 | 0.04 | 2.24 |
| *Ctxn3* | 16.45 | 36.58 | 0.04 | 2.22 |
| *Gprc5c* | 21.11 | 45.74 | 0.02 | 2.17 |
| *LOC120100227* | 41.83 | 90.27 | 0.04 | 2.16 |
| *Rsph3* | 39.96 | 85.64 | 0.02 | 2.14 |
| *Samd4a* | 118.04 | 244.14 | 0.02 | 2.07 |
| *Nrap* | 14.29 | 29.42 | 0.04 | 2.06 |
| *Cplx3* | 73.15 | 149.61 | 0.02 | 2.05 |
| *RGD1560171* | 29.56 | 59.55 | 0.04 | 2.01 |
| *Dok7* | 39.89 | 78.33 | 0.01 | 1.96 |
| *Niban1* | 51.18 | 99.65 | 0.05 | 1.95 |
| *Zfpm2* | 40.31 | 77.41 | 0.05 | 1.92 |
| *Spon2* | 49.66 | 95.11 | 0.03 | 1.92 |
| *Slc38a6* | 73.20 | 139.68 | 0.01 | 1.91 |
| *Sertad3* | 21.15 | 40.25 | 0.04 | 1.90 |
| *Htra3* | 98.74 | 186.23 | 0.03 | 1.89 |
| *LOC102555033* | 25.83 | 47.86 | 0.03 | 1.85 |
| *Mcm2* | 57.61 | 105.02 | 0.02 | 1.82 |
| *Trim34* | 95.45 | 173.66 | 0.02 | 1.82 |
| *Dcn* | 89.73 | 161.61 | 0.04 | 1.80 |
| *Cyb561a3* | 53.83 | 96.59 | 0.02 | 1.79 |
| *Foxq1* | 38.62 | 67.83 | 0.05 | 1.76 |
| *Plce1* | 61.16 | 106.22 | 0.04 | 1.74 |
| *Cryzl1-ps1* | 56.37 | 96.36 | 0.02 | 1.71 |
| *Ncf1* | 102.28 | 173.38 | 0.05 | 1.70 |
| *Entrep1* | 141.67 | 237.32 | 0.02 | 1.68 |
| *Ctu1* | 97.48 | 162.70 | 0.03 | 1.67 |
| *Galns* | 58.43 | 96.62 | 0.05 | 1.65 |
| *Eif4ebp1* | 75.52 | 124.64 | 0.05 | 1.65 |
| *Hes1* | 102.41 | 167.38 | 0.01 | 1.63 |
| *Mfng* | 54.76 | 89.38 | 0.04 | 1.63 |
| *Pde12* | 88.11 | 142.86 | 0.04 | 1.62 |
| *Dipk1c* | 93.27 | 150.34 | 0.02 | 1.61 |
| *Gpr137b* | 109.37 | 174.26 | 0.01 | 1.59 |
| *Mrc2* | 163.47 | 252.59 | 0.03 | 1.55 |
| *Ak3* | 463.68 | 689.93 | 0.05 | 1.49 |
| *Crlf1* | 325.94 | 482.82 | 0.05 | 1.48 |
| *Egfl7* | 315.68 | 458.81 | 0.00 | 1.45 |
| *Arhgef26* | 228.95 | 332.71 | 0.05 | 1.45 |
| *Fcsk* | 233.18 | 338.52 | 0.04 | 1.45 |
| *Sox9* | 428.92 | 620.38 | 0.02 | 1.45 |
| *Pdk1* | 147.03 | 211.55 | 0.02 | 1.44 |
| *Zfand2b* | 238.77 | 341.71 | 0.03 | 1.43 |
| *Lhx2* | 434.71 | 621.10 | 0.03 | 1.43 |
| *Prrc1* | 177.66 | 252.32 | 0.05 | 1.42 |
| *Ldlr* | 327.78 | 465.00 | 0.01 | 1.42 |
| *Fdxr* | 127.02 | 178.99 | 0.04 | 1.41 |
| *Dnase1l1* | 109.09 | 153.34 | 0.01 | 1.41 |
| *Vwf* | 573.51 | 793.25 | 0.01 | 1.38 |
| *Kdm2b* | 267.27 | 369.54 | 0.04 | 1.38 |
| *Mxra8* | 107.92 | 148.78 | 0.03 | 1.38 |
| *Phospho1* | 163.26 | 222.80 | 0.02 | 1.36 |
| *Lrp10* | 404.82 | 548.05 | 0.01 | 1.35 |
| *Mvk* | 313.20 | 417.05 | 0.04 | 1.33 |
| *Aldh16a1* | 255.02 | 337.69 | 0.01 | 1.32 |
| *Lrrc40* | 399.12 | 523.42 | 0.05 | 1.31 |
| *Nbl1* | 785.31 | 1020.07 | 0.03 | 1.30 |
| *Bscl2* | 526.63 | 670.03 | 0.04 | 1.27 |
| *Podxl2* | 800.85 | 1008.46 | 0.04 | 1.26 |
| *Paqr7* | 553.47 | 691.30 | 0.05 | 1.25 |
| *Isoc1* | 476.95 | 593.70 | 0.02 | 1.24 |
| *Mgat1* | 823.26 | 1015.86 | 0.04 | 1.23 |
| *Wasl* | 1474.97 | 1219.20 | 0.04 | -1.21 |
| *Zc2hc1a* | 534.11 | 420.33 | 0.05 | -1.27 |
| *Rnft1* | 392.79 | 307.98 | 0.04 | -1.28 |
| *Ppfia2* | 541.72 | 420.43 | 0.04 | -1.29 |
| *Atf2* | 1530.03 | 1175.55 | 0.03 | -1.30 |
| *Rbm25* | 1189.97 | 899.36 | 0.04 | -1.32 |
| *Abhd13* | 369.53 | 278.86 | 0.04 | -1.33 |
| *Iws1* | 287.58 | 209.85 | 0.04 | -1.37 |
| *Dimt1* | 246.64 | 179.75 | 0.04 | -1.37 |
| *Zc3h8* | 253.80 | 184.29 | 0.02 | -1.38 |
| *Fgf14* | 443.88 | 316.50 | 0.04 | -1.40 |
| *Hps3* | 251.62 | 178.71 | 0.02 | -1.41 |
| *Tmem231* | 203.17 | 142.87 | 0.03 | -1.42 |
| *Cbr4* | 168.68 | 118.37 | 0.03 | -1.43 |
| *Wdr36* | 335.28 | 234.36 | 0.03 | -1.43 |
| *Rerg* | 507.44 | 351.82 | 0.04 | -1.44 |
| *Adcy4* | 98.47 | 66.91 | 0.05 | -1.47 |
| *Tbc1d8b* | 100.97 | 66.07 | 0.04 | -1.53 |
| *Cdh9* | 473.65 | 305.20 | 0.02 | -1.55 |
| *Gpr83* | 139.56 | 89.77 | 0.02 | -1.55 |
| *Slc28a2* | 139.99 | 89.64 | 0.02 | -1.56 |
| *Fibcd1* | 185.93 | 116.33 | 0.05 | -1.60 |
| *ENSRNOG00000067837* | 132.80 | 81.01 | 0.01 | -1.64 |
| *Nfkbie* | 705.37 | 427.45 | 0.04 | -1.65 |
| *Samd9* | 517.02 | 312.48 | 0.01 | -1.65 |
| *ENSRNOG00000064438* | 168.30 | 98.06 | 0.01 | -1.72 |
| *Fgd4* | 179.56 | 103.63 | 0.04 | -1.73 |
| *ENSRNOG00000067762* | 124.11 | 70.29 | 0.02 | -1.77 |
| *LOC103691005* | 145.27 | 82.01 | 0.01 | -1.77 |
| *Pcdhb5* | 106.30 | 56.98 | 0.02 | -1.87 |
| *Irag1* | 170.43 | 89.80 | 0.00 | -1.90 |
| *Sms-ps2* | 116.92 | 53.92 | 0.01 | -2.17 |
| *C2h1orf162* | 39.45 | 17.22 | 0.05 | -2.29 |
| *ENSRNOG00000063571* | 31.91 | 13.22 | 0.04 | -2.41 |
| *Tet1* | 25.19 | 9.99 | 0.03 | -2.52 |
| *ENSRNOG00000068880* | 40.96 | 15.65 | 0.04 | -2.62 |
| *Lacc1* | 15.60 | 4.03 | 0.03 | -3.87 |
| *AABR07026565.1* | 19.34 | 4.86 | 0.04 | -3.98 |
| *Foxp3* | 15.47 | 3.85 | 0.02 | -4.02 |
| *ENSRNOG00000062327* | 18.44 | 4.52 | 0.04 | -4.08 |
| *ENSRNOG00000068110* | 19.48 | 4.71 | 0.05 | -4.13 |
| *RGD1560124* | 17.62 | 4.21 | 0.04 | -4.18 |
| *ENSRNOG00000070750* | 16.95 | 3.88 | 0.03 | -4.37 |
| *AABR07061707.1* | 28.89 | 6.58 | 0.03 | -4.39 |
| *ENSRNOG00000065247* | 16.58 | 3.63 | 0.04 | -4.57 |
| *ENSRNOG00000069485* | 16.92 | 3.57 | 0.04 | -4.73 |
| *ENSRNOG00000068740* | 9.78 | 1.69 | 0.04 | -5.79 |
| *Slc17a4* | 13.56 | 2.22 | 0.02 | -6.11 |
| *AABR07017236.2* | 12.22 | 1.76 | 0.03 | -6.93 |
| *ENSRNOG00000066670* | 7.48 | 0.98 | 0.04 | -7.60 |
| *LOC120093361* | 22.19 | 1.63 | 0.03 | -13.65 |
| *ENSRNOG00000067564* | 7.52 | 0.48 | 0.03 | -15.71 |
| *Acsm5* | 102.28 | 5.65 | 0.00 | -18.09 |

**CHMA1004 5 mg/kg**

| **Gene symbol** | **Average norm read counts Ctrl** | **Average norm read counts C4 (5mg/kg)** | **P-value** | **Fold change** |
| --- | --- | --- | --- | --- |
| *Amy1* | 12.39 | 1.00 | 0.03 | 12.39 |
| *RGD1565989* | 1.15 | 11.53 | 0.02 | 10.02 |
| *ENSRNOG00000062505* | 1.21 | 11.37 | 0.03 | 9.43 |
| *Olr1877* | 1.42 | 9.48 | 0.01 | 6.69 |
| *ENSRNOG00000069895* | 1.67 | 10.21 | 0.04 | 6.12 |
| *Jsrp1* | 1.71 | 10.22 | 0.03 | 5.99 |
| *LOC120093369* | 3.40 | 19.58 | 0.02 | 5.76 |
| *Clca4l* | 1.94 | 11.10 | 0.04 | 5.72 |
| *Sostdc1* | 3.07 | 16.25 | 0.05 | 5.30 |
| *ENSRNOG00000063128* | 2.18 | 10.25 | 0.04 | 4.69 |
| *Tab1* | 2.95 | 13.06 | 0.03 | 4.43 |
| *Rpl39-ps10* | 2.15 | 8.78 | 0.04 | 4.08 |
| *LOC100362298* | 5.47 | 21.76 | 0.04 | 3.98 |
| *C1h10orf95* | 2.57 | 10.17 | 0.05 | 3.95 |
| *Aph1bl1* | 3.79 | 14.50 | 0.04 | 3.83 |
| *Trdn* | 3.85 | 14.08 | 0.05 | 3.66 |
| *AABR07044570.2* | 6.94 | 24.62 | 0.05 | 3.55 |
| *Tpte2* | 3.35 | 11.11 | 0.05 | 3.32 |
| *Misp3* | 7.15 | 23.38 | 0.04 | 3.27 |
| *ENSRNOG00000063760* | 4.85 | 15.80 | 0.05 | 3.26 |
| *Tnfrsf26* | 3.66 | 11.93 | 0.03 | 3.26 |
| *ENSRNOG00000070983* | 3.82 | 12.13 | 0.03 | 3.17 |
| *Pou2af3* | 4.56 | 14.07 | 0.04 | 3.08 |
| *LOC108353164* | 4.86 | 14.80 | 0.04 | 3.05 |
| *LOC100912336* | 6.40 | 19.42 | 0.04 | 3.03 |
| *LOC102549694* | 46.32 | 140.30 | 0.01 | 3.03 |
| *Strc* | 6.37 | 18.53 | 0.03 | 2.91 |
| *ENSRNOG00000067969* | 5.45 | 15.46 | 0.02 | 2.84 |
| *RGD1561558* | 8.13 | 22.93 | 0.02 | 2.82 |
| *LOC100910341* | 4.55 | 12.73 | 0.05 | 2.80 |
| *Kif18a* | 6.46 | 17.97 | 0.03 | 2.78 |
| *LOC100360750* | 20.43 | 55.50 | 0.02 | 2.72 |
| *LOC100360828* | 13.79 | 36.52 | 0.02 | 2.65 |
| *AABR07044460.2* | 27.39 | 69.23 | 0.04 | 2.53 |
| *Tlr9* | 5.75 | 14.53 | 0.03 | 2.53 |
| *ENSRNOG00000067970* | 12.77 | 29.74 | 0.04 | 2.33 |
| *Evpl* | 12.52 | 28.95 | 0.01 | 2.31 |
| *M6pr-ps1* | 9.60 | 21.44 | 0.04 | 2.23 |
| *ENSRNOG00000070379* | 15.06 | 32.44 | 0.04 | 2.15 |
| *Zic4* | 17.81 | 36.09 | 0.02 | 2.03 |
| *Gramd1c* | 13.35 | 26.67 | 0.04 | 2.00 |
| *Car10* | 80.70 | 160.06 | 0.02 | 1.98 |
| *Serpinf2* | 27.06 | 53.19 | 0.01 | 1.97 |
| *Ppidl1* | 17.20 | 32.85 | 0.02 | 1.91 |
| *Rpl10l* | 31.65 | 60.24 | 0.04 | 1.90 |
| *Slc30a7* | 84.84 | 150.97 | 0.01 | 1.78 |
| *ENSRNOG00000019682* | 56.70 | 100.78 | 0.02 | 1.78 |
| *Fanca* | 25.98 | 45.39 | 0.02 | 1.75 |
| *Nxph3* | 186.82 | 317.88 | 0.02 | 1.70 |
| *Spc24* | 40.27 | 68.02 | 0.03 | 1.69 |
| *Mdga1* | 91.16 | 147.21 | 0.02 | 1.61 |
| *Nkain1* | 95.52 | 152.72 | 0.00 | 1.60 |
| *Fgf2* | 38.48 | 61.24 | 0.04 | 1.59 |
| *AABR07027854.1* | 58.63 | 92.23 | 0.01 | 1.57 |
| *Galns* | 61.85 | 96.82 | 0.04 | 1.57 |
| *Spon2* | 52.54 | 81.18 | 0.04 | 1.55 |
| *Dock8* | 88.01 | 135.02 | 0.02 | 1.53 |
| *Tdp1* | 61.72 | 92.60 | 0.02 | 1.50 |
| *Tmem255b* | 134.96 | 198.11 | 0.05 | 1.47 |
| *Ppl* | 61.02 | 88.60 | 0.04 | 1.45 |
| *Cpt2* | 111.95 | 158.15 | 0.03 | 1.41 |
| *Samd4a* | 124.90 | 174.88 | 0.04 | 1.40 |
| *Mocs1* | 174.58 | 244.21 | 0.04 | 1.40 |
| *Rhobtb3* | 114.31 | 158.41 | 0.03 | 1.39 |
| *Afg3l1* | 271.64 | 372.80 | 0.04 | 1.37 |
| *Sema3c* | 127.44 | 174.74 | 0.04 | 1.37 |
| *Rfk* | 245.48 | 335.18 | 0.04 | 1.37 |
| *Nbl1* | 830.93 | 1120.74 | 0.01 | 1.35 |
| *Synj2* | 275.42 | 370.26 | 0.04 | 1.34 |
| *Zswim9* | 112.71 | 147.20 | 0.05 | 1.31 |
| *Tmem186* | 250.71 | 314.11 | 0.03 | 1.25 |
| *Tfe3* | 355.37 | 440.53 | 0.04 | 1.24 |
| *Slc25a38* | 316.78 | 385.38 | 0.05 | 1.22 |
| *Asxl1* | 722.74 | 874.16 | 0.03 | 1.21 |
| *Arhgap12* | 491.33 | 400.11 | 0.04 | -1.23 |
| *Plrg1* | 426.58 | 339.33 | 0.03 | -1.26 |
| *Wee1* | 475.47 | 375.51 | 0.03 | -1.27 |
| *Gnpda2* | 323.43 | 249.21 | 0.04 | -1.30 |
| *Cfb* | 270.06 | 205.86 | 0.02 | -1.31 |
| *Rwdd4* | 458.91 | 348.91 | 0.01 | -1.32 |
| *Hps3* | 266.29 | 199.18 | 0.04 | -1.34 |
| *Ift74* | 206.15 | 153.20 | 0.02 | -1.35 |
| *Zfp35* | 252.33 | 185.34 | 0.02 | -1.36 |
| *Nme7* | 238.07 | 174.15 | 0.05 | -1.37 |
| *Filip1* | 417.20 | 301.56 | 0.03 | -1.38 |
| *Calr4* | 131.42 | 94.14 | 0.04 | -1.40 |
| *Vwa5b2* | 888.02 | 631.66 | 0.00 | -1.41 |
| *Col1a1* | 132.41 | 93.99 | 0.02 | -1.41 |
| *LOC102550944* | 132.37 | 93.82 | 0.03 | -1.41 |
| *Cbr1* | 443.47 | 309.99 | 0.04 | -1.43 |
| *Shbg* | 202.72 | 141.35 | 0.02 | -1.43 |
| *Bbof1* | 170.04 | 118.36 | 0.02 | -1.44 |
| *AABR07059663.1* | 143.94 | 97.01 | 0.02 | -1.48 |
| *Acta2* | 254.30 | 168.78 | 0.00 | -1.51 |
| *Peli1* | 691.61 | 456.42 | 0.00 | -1.52 |
| *Slc28a2* | 148.17 | 96.18 | 0.01 | -1.54 |
| *Dnaaf2* | 126.88 | 80.76 | 0.05 | -1.57 |
| *AABR07026797.1* | 54.96 | 34.45 | 0.03 | -1.60 |
| *Large2* | 53.65 | 31.73 | 0.03 | -1.69 |
| *ENSRNOG00000066365* | 183.98 | 107.45 | 0.02 | -1.71 |
| *Mx2* | 285.05 | 165.13 | 0.01 | -1.73 |
| *ENSRNOG00000065346* | 44.78 | 25.71 | 0.04 | -1.74 |
| *Abcg3l1* | 34.87 | 19.96 | 0.05 | -1.75 |
| *Ghdc* | 102.63 | 57.40 | 0.01 | -1.79 |
| *Il12a* | 44.65 | 24.45 | 0.04 | -1.83 |
| *Dnai3* | 41.30 | 21.81 | 0.05 | -1.89 |
| *LOC120099441* | 31.91 | 15.39 | 0.01 | -2.07 |
| *ENSRNOG00000056855* | 120.96 | 57.12 | 0.03 | -2.12 |
| *Lrrc32* | 52.02 | 23.72 | 0.01 | -2.19 |
| *LOC252890* | 13.27 | 3.07 | 0.04 | -4.32 |
| *AC112866.1* | 8.13 | 1.77 | 0.04 | -4.59 |
| *LOC120093339* | 82.65 | 16.20 | 0.00 | -5.10 |
| *ENSRNOG00000066670* | 7.92 | 1.51 | 0.04 | -5.25 |
| *RGD1563606* | 7.37 | 1.29 | 0.04 | -5.74 |
| *Pdilt* | 11.60 | 1.79 | 0.04 | -6.47 |
| *Ttc29* | 8.21 | 1.27 | 0.02 | -6.48 |
| *LOC120100339* | 6.37 | 0.75 | 0.03 | -8.44 |
| *Msx3* | 5.22 | 0.50 | 0.04 | -10.38 |
| *LOC681303* | 6.89 | 0.50 | 0.04 | -13.68 |
| *LOC120093361* | 23.51 | 0.77 | 0.02 | -30.33 |

**Common genes**

| **Gene symbol** | **C4 (1mg/kg) fold change** | **C4 (5mg/kg) fold change** |
| --- | --- | --- |
| *Misp3* | 2.92 | 3.27 |
| *Galns* | 1.65 | 1.57 |
| *Spon2* | 1.92 | 1.55 |
| *Samd4a* | 2.07 | 1.40 |
| *Nbl1* | 1.30 | 1.35 |
| *Hps3* | -1.41 | -1.34 |
| *Slc28a2* | -1.56 | -1.54 |
| *ENSRNOG00000066670* | -7.60 | -5.25 |
| *LOC120093361* | -13.65 | -30.33 |
