## Supplementary Table 2 for "Effect of RNA m^6^A methyltransferase activation by a low molecular weight compound on anxiety- and depression-related behaviours, monoamine neurochemistry and striatal gene expression in the rat"

**Supplementary Table 2.** The Database for Annotation, Visualization and Integrated Discovery (DAVID) gene ontology results based on differentially expressed genes (p<0.05) determined by RNA-Seq.

**CHMA1004 1 mg/kg**

| **GO term: Biological process** | **p-value** | **Genes** |
| --- | --- | --- |
| Phospholipase C-activating G-protein coupled receptor signaling pathway | 0.009 | *Oprd1, Plce1, Nmur2, Gpr83* |
| Positive regulation of phosphatidylinositol 3-kinase signaling | 0.014 | *Fgr, Ncf1, Sox9, Dcn* |
| Liver development | 0.017 | *Atf2, Vwf, Ak3, Hes1, Psm89* |
| Midbrain-hindbrain boundary morphogenesis | 0.025 | *Kdm2b, Hes1* |
| Cell adhesion | 0.029 | *Spon2, Egfl7, Vwf, Fibcd1, Mxra8, Pcdhb5, Hes1* |
| CD4-positive, CD25-positive, alpha-beta regulatory T cell differentiation | 0.03 | *Il2rg, Foxp3* |
| Gene expression | 0.032 | *Atf2, Ltb, Sox9, Il2rg, Foxp3* |
| Positive regulation of CD4-positive, CD25-positive, alpha-beta regulatory T cell differentiation | 0.042 | *Il2rg, Foxp3* |
| Negative regulation of smooth muscle contraction | 0.049 | *Ncf1, Irag1* |
| Bone mineralization | 0.05 | *Fgr, Phospho1, Sox9* |
| Response to bacterium | 0.053 | *Baiap2l1, Ncf1, Wasl, Mirlet7b, Psmb9* |
| Positive regulation of male gonad development | 0.055 | *Sox9, Zfpm2* |
| Lymphocyte proliferation | 0.067 | *Hells, Foxp3* |
| Apoptotic process | 0.069 | *Bard1, Hells, Atf2, Ncf1, Cdk1, Sox9, Mcm2* |
| Lipid metabolic process | 0.072 | *Bscl2, Acsm5, Lrp10, Ldlr* |
| Protein localization to kinetochore | 0.072 | *Mtbp, Cdk1* |

| **GO term: Cellular component** | **p-value** | **Genes** |
| --- | --- | --- |
| Mitochondrial matrix | 0.002 | *Pde12, Dimt1, Cdk1, Ak3, Acsm5, Cbr4, Pdk1* |
| Receptor complex | 0.054 | *Amn, Il2rg, Ldlr, Gprc5c, Crlf1* |

| **GO term: Molecular function** | **p-value** | **Genes** |
| --- | --- | --- |
| Metal ion binding | 0.01 | *Bard1, Atf2, Spon2, Loc103691005, Loc102555033, Adcy4, Zc3h8, Acsm5, Cyb561a3, Fop3, Asrg1, Rnft1, Galns, Fgd4, Phospho1, Lhx2, Pde12, Mfng, Nrap, Plce1, Zfpm2, A3galt2, Mcm2* |
| Oxidoreductase activity | 0.027 | *Fdxr, Rdh16, Aldh16a1, Cyb561a3, Cbr4* |
| ATP binding | 0.051 | *Rgd1561440, Polq, Hells, Mvk, Adcy4, Ak3, Acsm5, Fign, Fgr, Fcsk, Myh8, Cdk1, Kif20b, Pdk1, Mcm2* |
| Low-density lipoprotein receptor activity | 0.063 | *Lrp10, Ldlr* |
| Translation repressor activity | 0.063 | *Samd4a, Eif4ebp1* |
| Collagen binding | 0.068 | *Mrc2, Vwf, Dcn* |
| Identical protein binding | 0.091 | *Polq, Atf2, Mvk, Vwf, Htra3, Ak3, Wasl, Foxp3, Cdh9, Asgr1, Prrc1, Rdh16, Hes1, Nbl1, Ldlr, Pdk1, Trim34* |
| Actin binding | 0.091 | *Fgd4, Baiap2l1, Nrap, Myh8, Wasl* |
| Neurotransmitter transporter activity | 0.091 | *Slc28a2, Cplx3* |

| **KEGG pathway** | **p-value** | **Genes** |
| --- | --- | --- |
| Chemical carcinogenesis – receptor activation | 0.053 | *Atf2, Paqr7, Eif4ebp1, Adcy4, Mirlet7b* |
| Cortisol synthesis and secretion | 0.067 | *Atf2, Adcy4, Ldlr* |
| Thyroid hormone synthesis | 0.076 | *Atf2, Adcy4, Asgr1* |
| cGMP-PKG signaling pathway | 0.091 | *Oprd1, Atf2, Irag1, Adcy4* |

**CHMA1004 5mg/kg**

| **GO term: Biological process** | **p-value** | **Genes** |
| --- | --- | --- |
| Response to virus | 0.006 | *Acta2, Mx2, Tlr9, Il12a* |
| Regulation of antibacterial peptide production | 0.012 | *Ppl, Evpl* |
| Wound healing | 0.024 | *Col1a1, Ppl, Evpl, Fgf2* |
| Response to lipopolysaccharide | 0.029 | *Spon2, Cbr1, Peli1, Il12a, Cfb* |
| Inner dynein arm assembly | 0.059 | *Dnai3, Dnaaf2* |
| Intermediate filament cytoskeleton | 0.087 | *Ppl, Evpl* |
| Animal organ morphogenesis | 0.099 | *Asxl1, Nbl1, Fgf2* |

| **GO term: Cellular component** | **p-value** | **Genes** |
| --- | --- | --- |
| Cytoplasm | 0.026 | *Ghdc, Dock8, Rhobtb3, Loc100360750, Ttc39, Evpl, Ppl, Ppidl1, Fgf2, Arhgap12, Il12a, Tpte2, Slc30a7, Cbr1, Dnai3, Dnaaf2, Mx2, Tfe3, Samd4a, Hps3, Synj2, Col1a1, Acta2, Ac112866.1, Wee1, Kif18a, Pou2af3, Gnpda2, Nme7, Tdp1, Tlr9, Filip1, Bbof1* |
| Extracellular space | 0.075 | *Col1a1, Spon2, Sema3c, Serpinf2, Sostdc1, Il12a, Amy1, Shbg, Lrrc32, Nbl1, Fgf2, Cfb* |
| Cytoskeleton | 0.099 | *Acta2, Rhobtb3, Ttc29, Ppl, Evpl* |

| **GO term: Molecular function** | **p-value** | **Genes** |
| --- | --- | --- |
| Translation repressor activity | 0.067 | *Ac112866.1, Samd4a* |
| Bmp binding | 0.078 | *Sostdc1, Nbl1* |

| **KEGG pathway** | **p-value** | **Genes** |
| --- | --- | --- |
| Toll-like receptor signalling pathway | 0.054 | *Tlr9, Il12a, Tab1* |
| Chagas disease | 0.066 | *Tlr9, Il12a, Calr4* |
